# Biological context modulates virus-host dynamics and diversification

**DOI:** 10.1101/2025.09.10.675297

**Authors:** Rodrigo Sanchez-Martinez, Esther Rubio-Portillo, Laura Medina-Ruiz, Jonás Sarasa, María Enciso, Fernando Santos, Josefa Antón

**Affiliations:** Department of Physiology, Genetics and Microbiology, University of Alicante, San Vicent del Raspeig, Alicante, Spain; IGLS, Alicante, Spain; Institute of Health and Biomedical Research of Alicante (ISABIAL), Alicante, Spain; Multidisciplinary Institute of Environmental Studies Ramon Margalef, Alicante, Spain

## Abstract

Virus-host interactions are fundamental to microbial ecology and evolution, yet they are often studied in simplified systems. To assess the impact of biological complexity, we examined the halophilic bacterium *Salinibacter ruber* strain M1 and the EM1 virus, a bacteriophage that infects M1, in the presence of additional strains and viruses. In the short term, the presence of a biotic background delayed population-level lysis and reduced EM1 production. In the long term, both M1 and EM1 persisted across all conditions, but their evolutionary trajectories diverged. The virus accumulated many mutations, showed reduced infectivity against its native host, and expanded its host range in the presence of other viruses, highlighting a role of virus-virus interactions in promoting diversification. In contrast, the host accumulated more mutations when evolving with other bacterial strains. Taken together, these findings demonstrate that biological complexity shapes both the ecological dynamics and evolutionary trajectories of virus-host interactions.

## INTRODUCTION

Natural ecosystems are highly complex networks where multiple biotic and abiotic factors interact dynamically (Fuhrman et al., 2015; Levin, 1998). This complexity makes it challenging to predict ecological and evolutionary phenomena since experimental studies are generally conducted under simplified laboratory conditions which do not reflect natural dynamics (Calisi & Bentley, 2009; Chen et al., 2023; Roberts et al., 2015; Zimmer & Dorea, 2023). To advance our understanding of microbial processes and predict how microbial communities respond to environmental changes or selective pressures, it is crucial to design experiments that more faithfully simulate natural conditions, incorporating the diversity of species and interactions that characterize real ecosystems (Benton et al., 2007; Blazanin & Turner, 2021; Castledine & Buckling, 2024; Jessup et al., 2004).

Viruses act as major eco-evolutionary forces in microbial ecosystems. As the most abundant biological entities on Earth, they play a central role in regulating microbial communities through diverse infection cycles that impact nutrient cycling, genetic diversity, and ecosystem structure (Fierer, 2017; Middelboe & Brussaard, 2017; Rohwer & Thurber, 2009). Virus-host interactions also drive microbial evolution, triggering adaptations that influence community composition and function. Despite their importance, many aspects of these interactions remain poorly understood, particularly under complex, natural conditions. Although numerous studies have explored virus-host interactions, most have relied on simplified experimental systems involving a single virus-host pair under controlled conditions, or on metagenomic approaches that capture complex communities but lack experimental resolution (Camarillo-Guerrero et al., 2021; Coutinho et al., 2017; Ghatbale et al., 2025; Neri et al., 2022; Scanlan et al., 2015; Scanlan & Buckling, 2012). As a result, we still need an understanding of how community context, particularly the simultaneous presence of multiple hosts and viruses, shapes infection dynamics and viral evolution. In addition, recent works suggest that viruses can influence microbial communities at the strain level rather than at the species level, highlighting the importance of considering intraspecific diversity when studying these interactions (Castledine & Buckling, 2024; Goyal et al., 2022). Bridging this gap requires experimental systems that incorporate intermediate levels of biological complexity while maintaining tractability. Some recent works have moved in this direction (Blazanin & Turner, 2021; Castledine & Buckling, 2024; Chevallereau et al., 2022; Ferriol-González & Domingo-Calap, 2025; Koskella et al., 2022). However, these studies did not explicitly address how multiple co-occurring viruses interact, nor how these interactions unfold over longer evolutionary timescales. This limitation hinders our ability to interpret patterns observed in natural microbial communities and to predict virus-host dynamics in complex ecosystems.

Hypersaline aquatic environments, which constitute approximately half of continental waters, harbor microbial communities that, at extreme salinities, are dominated by archaea (Burns et al., 2007; Shiklomanov, 1998; Ventosa, 2006). At these salinities, *Salinibacter ruber* emerges as the main bacterial species (Antón et al., 2002; Gomariz et al., 2015; Oren, 2011). Although *Sal. ruber* is as halophilic as the dominant archaea at close-to-saturation salinities, it is consistently less abundant, representing between 1% and 10% of total cells, indicating that it could be subjected to intense selective pressure (Mora-Ruiz et al., 2018; Ventosa et al., 2015). Cultivation and metagenomic studies reveal a high degree of intraspecific genomic diversity in *Sal. ruber*, including the coexistence of distinct phylogroups and adaptation to local microenvironments (Viver et al., 2015, 2023). These hypersaline habitats also harbor some of the highest recorded viral concentrations, with virus-like particles (VLP) reaching up to 10^10^ VLP/ml and virus-to-cell ratios ranging from 10 to 100 (Di Meglio et al., 2016; Santos et al., 2012). Given its environmental ubiquity, cultivability, and intraspecific variation, likely shaped in part by viral predation, *Sal. ruber* constitutes a valuable model for studying microbial ecology and virus-host interactions.

In this work, we have addressed how the ecology and evolution of a virus-host pair (i.e., *Sal. ruber* strain M1 and the EM1 virus) are affected by the presence of additional *Sal. ruber* strains and their viruses, using an experimental system that captures intermediate levels of biological complexity. By increasing community complexity, we have demonstrated that: (i) the presence of other strains and viruses delays viral production and reduces EM1 virion production; and that (ii) the biological context influences the long-term evolution of the virus-host pair and promotes changes in viral infectivity and host range, driven by a higher number of mutations under more complex contexts. These findings highlight the importance of considering biological complexity in studies of virus-host interactions, considering both host and viral diversity essential to better understand and predict microbial eco-evolutionary processes in natural system. Our work not only contributes to a better understanding of these dynamics in natural systems but also suggests that this complexity should be considered in practical applications.

## RESULTS

Five parallel experiments, with biological triplicates each, evaluated the effects of the presence of *Sal. ruber* strains and viruses on the interactions and evolution of *Sal. ruber* strain M1 and the EM1 virus, a bacteriophage that specifically infects strain M1 (Villamor et al., 2018) (Fig. 1a, b). In experiment 1, *Sal. ruber* strain M1 was cultured alone. In experiment 2, *Sal. ruber* strains M8, M31, and P18 were added to the M1 culture. The bacterial strains used represent closely related members of the *Sal. ruber* population, capturing natural intraspecific diversity. Genome comparisons previously indicated high average nucleotide identity (ANI) values among strains (typically >99%) (González-Torres & Gabaldón, 2018), consistent with strain-level variation within environmental populations, while still maintaining distinct phenotypic and ecological traits. Several of these strains show high similarity to isolates recovered from geographically distant hypersaline environments more than 20 years after its first isolation, supporting their ecological relevance and potential coexistence in nature (Conrad et al., 2022, 2024; Viver et al., 2024). Experiments 1 and 2 served as virus-free controls.

**Fig. 1:**
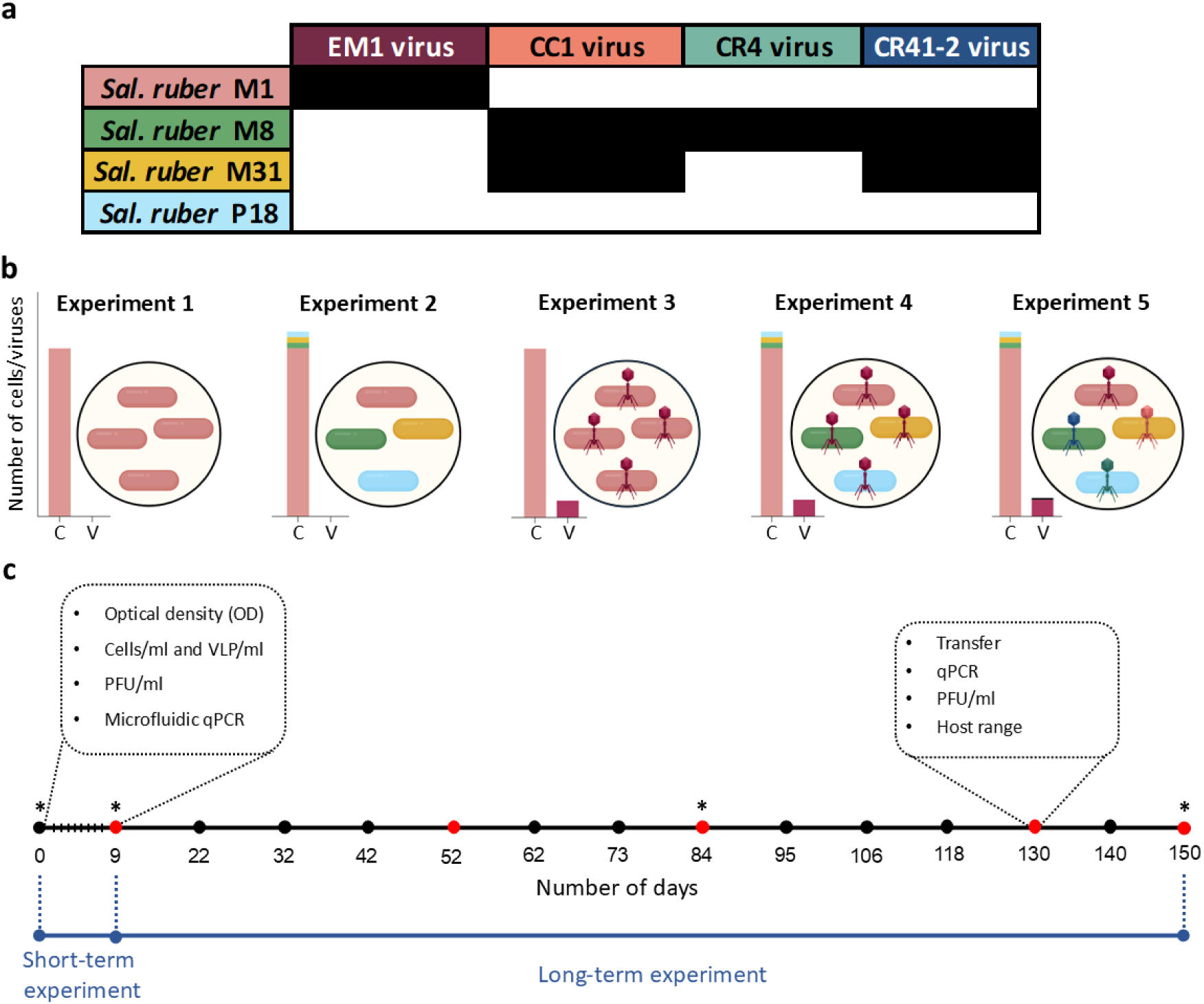
Experimental design. **a)** Host range of the viruses used in this study. Viruses are indicated at the top, and *Sal. ruber* strains on the left. Black and white rectangles indicate infection and no infection, respectively. **b)** Overview of the five experiments using *Sal. ruber* M1. Experiments 2, 4, and 5 additionally included strains M8, M31, and P18. Experiments 3, 4, and 5 included the EM1 virus, and experiment 5 also included the CC1, CR41-2, and CR4 viruses. The color bars next to each experiment indicate the proportion of each strain (cells/ml, left bars) or virus (VLP/ml, right bars). All experiments were performed in biological triplicates. **c)** Timeline of the short- and long-term experiments. Short-term samples were incubated for 9 days with regular monitoring. Long-term experiments were transferred to fresh medium every 9–13 days over 150 days (black and red dots), with additional parameters measured at selected time points (red dots). Extracellular viral assemblages from experiments 3, 4, and 5 were sequenced at 9, 84, and 150 days, along with the original viral stocks (asterisks indicate points chosen for sequencing). Figure created with Biorender.

In experiment 3, M1 alone was infected with its EM1 virus. Experiment 4 included M1 infected with EM1, along with the presence of strains M8, M31, and P18. Finally, experiment 5 included the same host as experiment 4, plus M31CC1 and M31CR41-2 viruses (referred to as CC1 and CR41-2, respectively), which infect both *Sal. ruber* strains M31 and M8, and M8CR-4 virus (i.e. CR4), which infects *Sal. ruber* strain M8 (Villamor et al., 2018) (Fig. 1a). The viruses, none of them able to infect M1, were selected to represent different ecological interaction patterns within the *Sal. ruber* community. Specifically, we added viruses with different host-ranges, comprising more generalists, capable of infecting multiple strains (CC1 and CR41-2), as well as a virus with a narrower host range (CR4). We also incorporated a bacterial strain (P18) that is not infected by any of the viruses used in the experiment. The selected viruses belong to three different genera (Supplementary Fig. 1; Villamor et al., 2018) and display distinct life-history traits, with three virulent viruses and one with potential lysogenic capability (CC1, which encodes an integrase). These viruses have been detected in multiple environmental viromes more than a decade after their isolation (data not published), suggesting that they can coexist in natural hypersaline systems over extended periods.

The first 9 days of incubation (“short-term experiment”), corresponding to ∼14 generations of *Sal. ruber* (with a generation time ∼15 hours), evaluated infection traits such as lysis timing, virion production, infectivity (fraction of viral particles capable of initiating productive infections on the original host), and community dynamics. Thereafter, all cultures were periodically transferred to fresh medium and maintained for 150 days (∼240 generations). This constituted the “long-term experiment”, aimed at studying possible evolutionary changes in the virus-host pair, while continuing to monitor key parameters of the infection (Fig. 1c).

### Biological complexity delays population-level lysis

In the short-term experiment, *Sal. ruber* exhibited its characteristic growth pattern both in monoculture and in the presence of other strains, with a slightly higher optical density (OD) in experiment 2, likely due to the contribution of the additional bacterial strains (Fig. 2a; Supplementary Fig. 2a). In the infection experiment (experiment 3), lysis of *Sal. ruber* M1 was observed, followed by a recovery, a pattern that mimics previous findings from our laboratory, in which that growth was mainly attributed to the emergence of pseudolysogens (Sanchez-Martinez et al., 2025). In the experiments which contained other strains (4 and 5), this OD recovery was observed earlier, which could be attributed to the growth of uninfected strains.

**Fig. 2:**
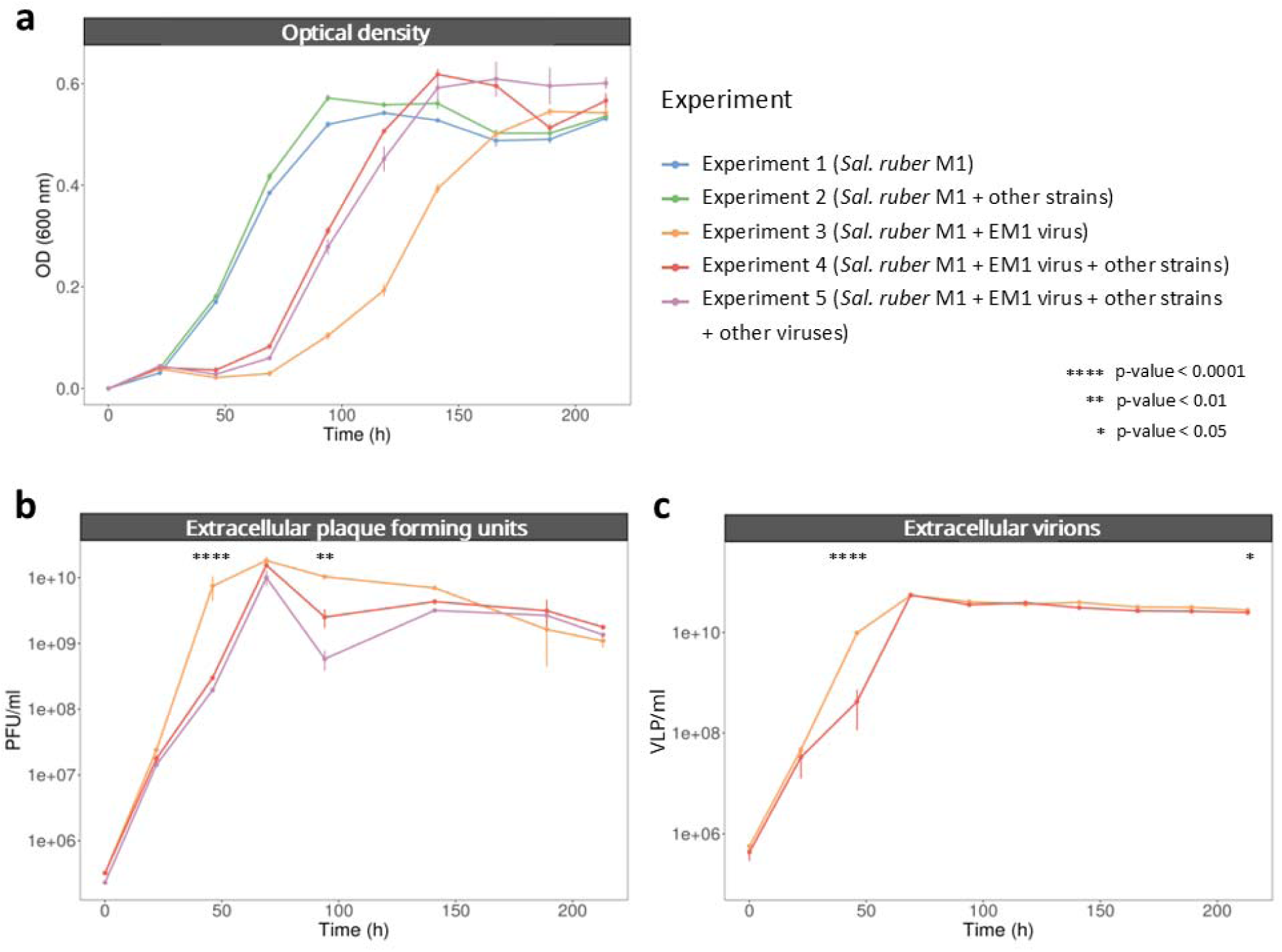
Influence of the biological context on the virus-host dynamics in the *Sal. ruber* M1-EM1 pair. **a)** Optical density at 600 nm of all five short-term experiments. **b)** PFU/ml of experiments 3, 4 and 5, measured by plaque assay against the original host *Sal. ruber* M1. **c)** Extracellular VLP/ml of experiments 3 and 4 measured by flow cytometry. All experiments were performed in biological triplicates. Error bars represent the standard error of the mean across replicates. Statistical significance was assessed using pairwise t-tests between experiment 3 and experiments 4 and 5, with p-values indicated by asterisks. ‘Other strain’ refers to M8, M31 and P18 strains and ‘other viruse’ refers to CC1, CR4, and CR41-2 viruses.

The production of infectious EM1 viral particles, measured as plaque-forming-units (PFU) on *Sal. ruber* M1, was significantly delayed at 48 hours in experiments including other strains (experiment 4) and both additional strains and viruses (experiment 5) compared to the virus-host pair alone (*t*-test p-value < 0.0001) (Fig. 2b). This pattern indicated a delay in M1 population-level lysis dynamics and viral production in the presence of additional *Sal. ruber* strains. After 96 hours, a drop in the number of infectious particles was observed, more pronounced in the more complex experiments (p-value < 0.01). In all the experiments a slight decline in the number of PFUs was observed over time, with no significant differences between days 6 and 9. Total extracellular virion concentration, measured by flow cytometry, reached 9.7 x 10^10^ VLP/ml within 48 hours in absence of the biological background, whereas in the experiment with other strains of *Sal. ruber* this concentration was 4.2 x 10^8^ VLP/ml, a statistically significant reduction (p-value < 0.0001) (Fig. 2c), consistent with the observations of PFU counts. The more complex experiment was excluded from this figure, as the presence of multiple coexisting viruses impedes attributing VLP counts specifically to EM1 (Supplementary Fig. 2b).

High throughput microfluidics-based qPCR revealed that the concentration of the M1 strain (measured as genomes/ml) varied across experiments and time points (Fig. 3). At 48 hours, M1 abundance dropped significantly in the treatment where only the M1-EM1 pair was present (3.9 x 10^6^ genomes/ml), indicating active lysis of the population, whereas it remained much higher in the more complex conditions including additional strains (1.2 x 10^9^ genomes/ml) or both additional strains and viruses (7.6 x 10^8^ genomes/ml) (p-value < 0.0001). By 72 hours, M1 levels decreased in all experiments, maintaining low values in the simple infection (3.8 x 10^6^ genomes/ml), and showing delayed but evident reductions in the experiments including additional strains and both additional strains and viruses (2.2 x 10^6^ and 5.2 x 10^6^ genomes/ml, respectively) (Fig. 3; Supplementary Fig. 3).

**Fig. 3:**
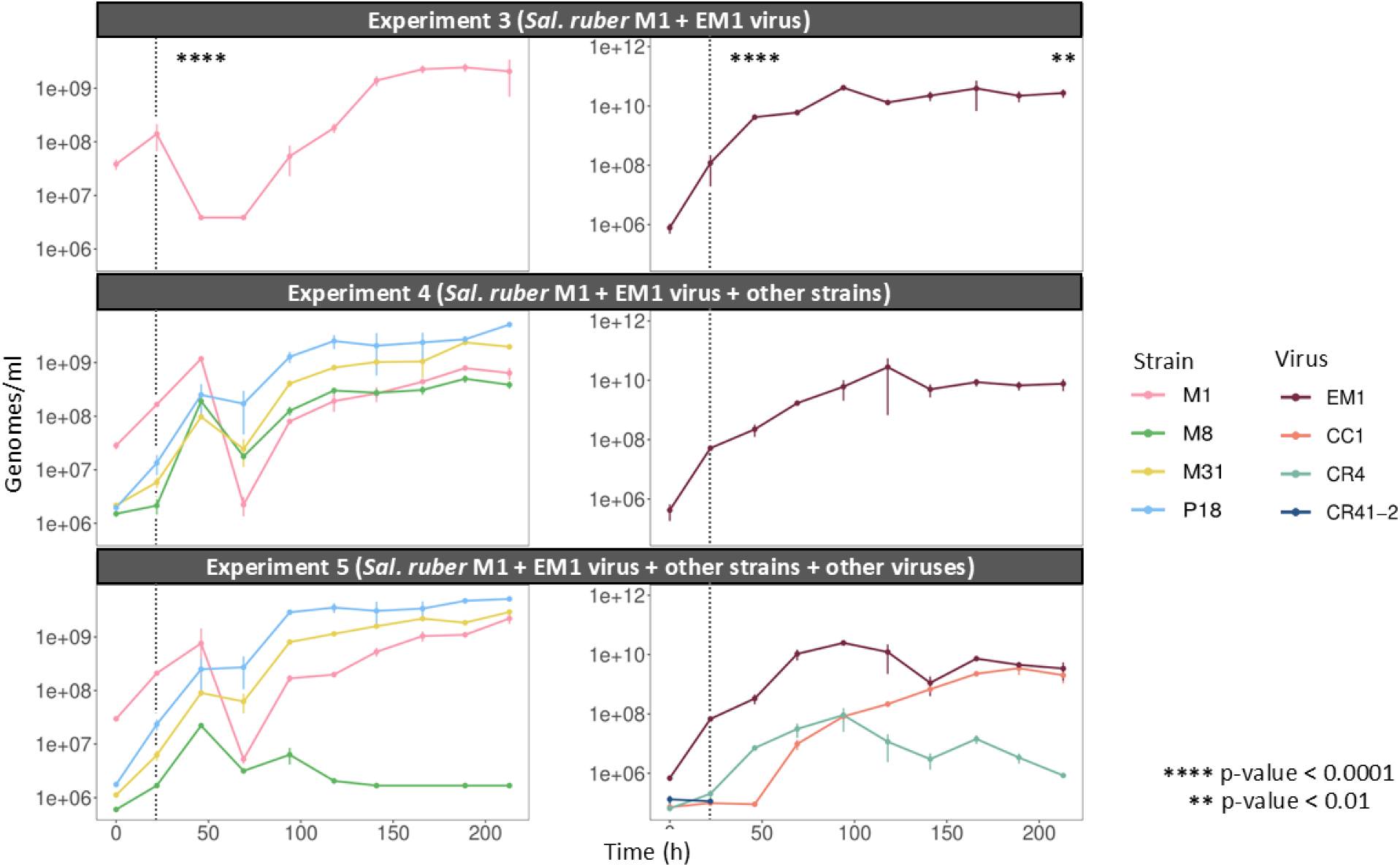
Concentration of the different *Sal. ruber* strains and viruses in experiments 3, 4, and 5. Genomes/ml of each bacterial strain in experiments 3, 4 and 5 (top to bottom) measured by microfluidic qPCR (left). Virus free genomes/ml in the supernatant of experiments 3, 4, and 5 (top to bottom) measured by microfluidics-based qPCR (left). All experiments were performed in triplicate. Error bars represent the standard error of the mean across replicates. Statistical significance was assessed using pairwise t-tests between experiment 3 and experiments 4 and 5, with p-values indicated by asterisks.

Regarding the bacterial community, in the infection experiment including additional strains (experiment 4), a decrease in the concentration of M8, M31, and P18 strains was observed at 50 hours, despite the absence of their viruses (Fig. 3). This pattern suggests that the growth of these strains may be transiently inhibited after *Sal. ruber* M1 lysis. To test this, a virus-free lysate from an M1-infected culture was added to pure cultures of M8, M31, and P18. A clear, transient growth inhibition was observed, confirming that the M1 lysate affected the growth of the other strains (p-value at 113 hours: < 0.01 for M8; < 0.01 for M31; <0.05 for P18) (Supplementary Fig. 4). In the more complex experiment, M8 showed minimal growth. As the only strain infected by three viruses (Fig. 1a), M8 likely could not withstand the high viral pressure, explaining its poor performance under these conditions.

When extracellular viruses were quantified by microfluidics-based qPCR, lower EM1 concentrations were observed at 48 hours in conditions including additional strains (2.2 × 10 genomes/ml) and both additional strains and viruses (3.3 × 10 genomes/ml), compared to the virus-host pair alone (4.2 x 10^9^ genomes/ml) (p-value < 0.0001) (Fig. 3; Supplementary Fig. 5), confirming that increased community complexity delayed the infection and lysis of *Sal. ruber* M1 by the EM1 virus. By 216 hours (day 9), experiment 4, including other strains, yielded a significantly lower VLP concentration than experiment without biological background (2.4 x 10^10^ and 2.8 x 10^10^ VLP/ml, respectively, p-value < 0.05) (Fig. 2c). Microfluidics-based qPCR measurements also revealed that the concentration of extracellular EM1 virus in experiments including additional strains and both additional strains and viruses (7.6 x 10^9^ and 3.4 x 10^9^ genomes/ml, respectively) was significantly lower than in experiment where only the M1-EM1 pair was present (2.7 x 10^10^ genomes/ml) at the same time point (p-value < 0.01), supporting the flow cytometry results (Fig. 3). These findings indicated that the presence of additional *Sal. ruber* strains reduced the overall production of EM1.

The more complex experiment, including all the strains and viruses, also revealed distinct patterns in the production of other viruses. CC1 virus, which infects both *Sal. ruber* M8 and M31, showed sustained high titers from 50 hours onward, consistent with ongoing replication in its hosts (Fig. 3). CR41-2 virus, which infects the same strains as CC1 (M8 and M31), was only detected on days 1 and 4. Adsorption assays revealed that CR41-2 adsorbs significantly less efficiently than CC1 to M31 (Supplementary Fig. 6). While differences in adsorption may contribute to these dynamics, other factors, such as latent period, burst size, or general population dynamics, could also play a role. Finally, CR4 virus, which exclusively infects M8, exhibited an early increase followed by a sharp decline (Fig. 3). This decline likely reflects the lack of M8 cell growth, which limited the availability of susceptible hosts and consequently reduced further CR4 replication.

### Host and virus establish a stable long-term relationship

After the initial 9-day period, all cultures were transferred to fresh medium to monitor their long-term evolution with transfers every 9-13 days over a total of 150 days. At selected time points, specific parameters were assessed, including changes in host range and infectivity toward the native host (Fig. 1c).

*Sal. ruber* M1 was maintained at high density (∼10 genomes/ml) across all time points, regardless of the presence of EM1 or other strains and viruses (Supplementary Fig. 7). The EM1 virus maintained relatively stable concentrations extracellularly across all three experiments (Supplementary Fig. 8; Supplementary Fig. 9). Although the EM1 titer declined by about one order of magnitude between days 52 and 84, its level remained stable until the end of the study. These results indicate that *Sal. ruber* M1 and its EM1 virus established a stable coexistence that persisted throughout the study, regardless of the surrounding biological context. Long-term virus-host coexistence has been widely reported in other systems driven by mechanisms such as arms-race dynamics, spatial heterogeneity, or phase variation in gene expression (Cortés-Martín et al., 2025; Kortright et al., 2022; Lourenço et al., 2020). However, a previous work from our group demonstrated that EM1 can be maintained in *Sal. ruber* M1 in a pseudolysogenic state (Sanchez-Martinez et al., 2025), which appears to underlie the stable coexistence observed here. To confirm this, colonies were isolated at the final time point (150 days). Nine colonies (three from each biological replicate) from experiment 3, where only the M1-EM1 pair was present, and nine from experiment 5, the more complex experiment, could be identified as M1 by PCR. A parallel PCR with EM1-specific primers showed that all colonies contained the viral genome, demonstrating that EM1 was maintained in a pseudolysogenic state. All colonies were also resistant to both the initial EM1 virus and the final virus assemblage, confirming the role of pseudolysogeny in long-term virus-host coexistence and in bacterial resistance (Supplementary Fig. 10). Importantly, these results indicate that the establishment and maintenance of EM1 in M1 occur independently of the presence of additional strains or viruses, highlighting the potential robustness of this interaction in more complex contexts.

Additional insights emerged from the dynamics of the other community members. In experiment 5, CC1 remained at high concentrations, as well as one of their hosts, M31, which was also detected until the end of the experiment (Supplementary Fig. 7 and 8). In the case of the CR4 virus, although it was not detected extracellularly after 9 days, it persisted in the total culture until 84 days, coinciding with the extinction of its host M8 (Supplementary Fig. 9). The P18 strain, the only one not infected by any virus, disappeared in experiment 5 from all 3 replicates and, although still detectable, was outcompeted by other strains in experiments 2 and 4. This indicates that viruses were not the only force structuring the community, and that competitive interactions among bacterial strains also played a crucial role.

### Biological context reduces viral infectivity and alters host range

To assess the infective capacity of the viral population, we estimated infectivity as the ratio between PFU/ml (measured using the native M1 strain as the host) and extracellular EM1 genomes/ml. This metric reflects the fraction of viral particles capable of initiating productive infections on the original host, rather than total viral abundance. On day 9, the infectivity of the EM1 virus was maintained above 0.1% in the three experiments and remained stable at these levels for 52 days (Fig. 4a). However, by day 84, significant differences (p-value < 0.0001) emerged between the experiment 3, where only the M1-EM1 pair was present, and those more complex (experiments 4 and 5). In the simplest infection (experiment 3), infectivity remained stable but, in experiment 4, where the *Sal. ruber* M1-EM1 pair was accompanied by other strains, infectivity dropped by almost one order of magnitude. In the most biologically complex (experiment 5), this reduction in infectivity was more drastic, with a drop of nearly 4 orders of magnitude compared to the simplest one. By day 130, infectivity had decreased further in all experiments, down to 5 orders of magnitude below the initial level. These findings suggested that the EM1 virus underwent genomic changes, as further explored below, which reduced its ability to infect the native strain in all the experiments and highlighted that the biological context significantly affected the interactions between EM1 and its host.

**Fig. 4:**
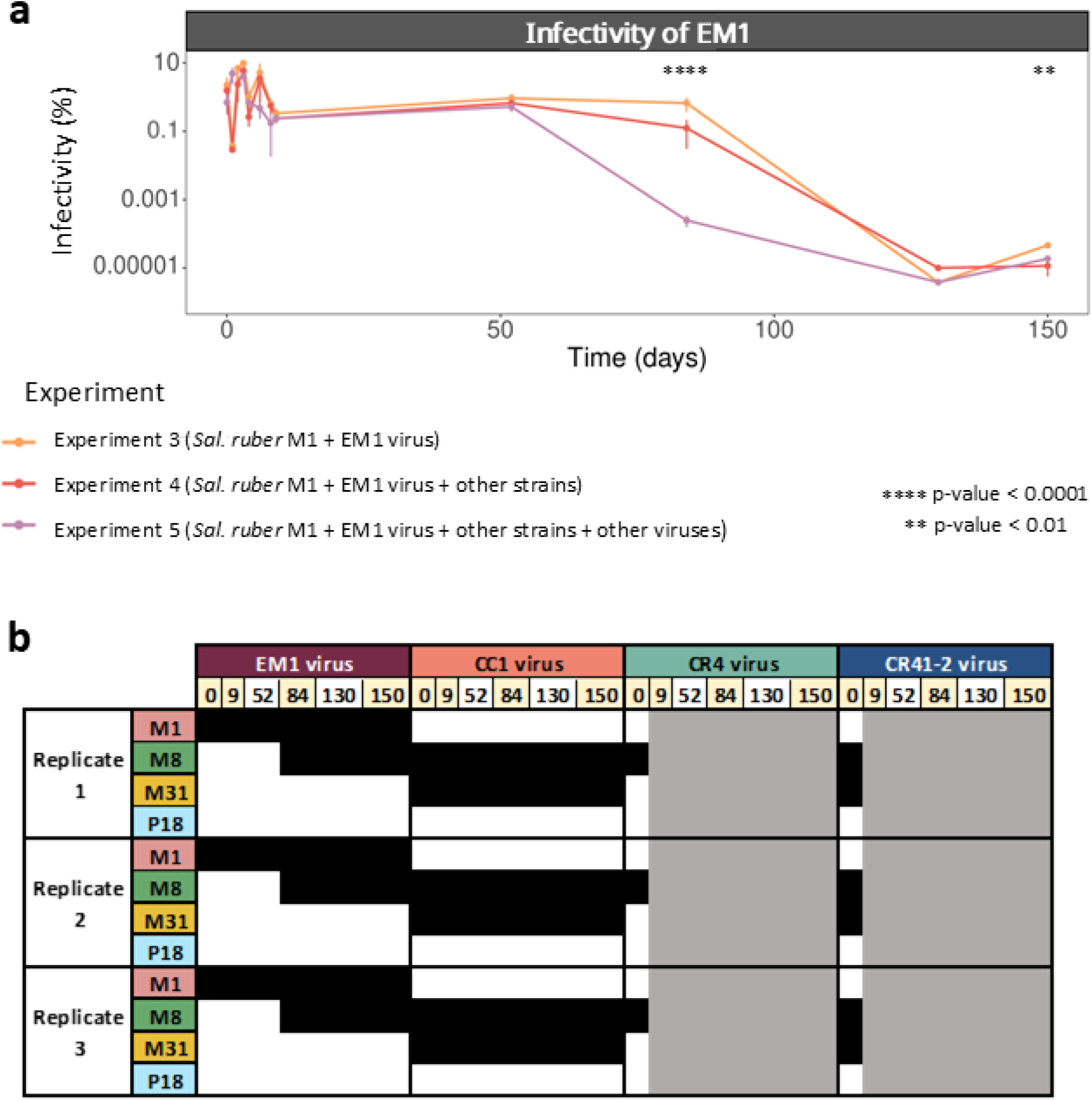
The biological context affects the evolution of the infectivity and host range of the EM1 virus. **a)** Infectivity of EM1 virus in experiments 3, 4 and 5 in the long-term experiment, defined as the ratio between PFU/ml and extracellular genomes/ml. The y-axis represents the percentage of infectious viruses. Error bars represent the standard error of the mean across replicates. Statistical significance was assessed using pairwise t-tests between experiment 3 and experiments 4 and 5, with p-values indicated by asterisks. **b)** Host range of the four viruse across the three biological replicates in experiment 5. Black squares indicate infection, and white squares indicate no infection. The gray area indicates no viral detection in the extracellular fraction. The time points marked in soft yellow (upper part) denote those selected for genomic analysis.

Host range changes were also detected in the EM1virus in the most complex experiment 5, whereas no such changes were observed in the experiments 3 and 4, where only the M1-EM1 pair was present and those including additional strains, respectively. The evolved EM1 in experiment 5, which initially only infected the M1 strain, expanded its host range and infected the native M8 strain consistently across all three biological replicates at 84 hours (Fig. 4b). These findings suggest that the observed host range expansion in EM1 was a result of selective pressures arising from increased ecological complexity. Coexistence with other strains and viruses may have favored viral variants capable of infecting alternative hosts (i.e., either by exploiting new receptor variants or by overcoming host-specific resistance mechanisms (see below)). In the case of the other viruses, none of them showed any host range changes.

The unequal decline in the EM1 virus infectivity across experiments, together with the shift in host range observed in the experiment involving other strains and viruses, suggested that EM1 underwent different genomic changes under the different conditions, influencing the virus-host evolutionary trajectory according to the surrounding biotic environment. To analyze whether genomic changes could be responsible for these differences, extracellular viral assemblages and cellular pellets from the experiments of the infections (3, 4 and 5) were sequenced on days 9, 84 and 150. The three biological replicates of each experiment were pooled prior to DNA extraction. Pooling was performed to capture the overall mutational landscape representative of each condition, with equal amounts of DNA from each replicate combined to ensure balanced representation, allowing the identification of mutations consistently present across replicates while minimizing replicate-specific variation. The original viral genomes were also sequenced as references.

### Viral genomic diversification is enhanced by other viruses

EM1 genome remained largely stable during the early stages of the experiment, with only a single mutation in a gene coding for a hypothetical protein detected across all treatments by day 9 (Supplementary Table 1). By day 84, the number of mutations remained low in the condition with only the M1-EM1 pair and in the condition including additional strains (15 and 14 mutations, respectively), mostly in non-coding regions (Supplementary Fig 11; Supplementary Table 1). In contrast, the condition including both additional strains and viruses (experiment 5) showed a markedly different pattern, with 772 mutations distributed across the genome. A large proportion of these occurred in structural genes, particularly those related to tail components. Two genes which code for minor tail proteins showed over 190 mutations, and a gene encoding a long-tail fiber protein, 69 mutations. This genomic diversification coincided with a sharp drop in EM1 infectivity against the native M1 host, pointing to a possible link between these mutations in tail genes and host range evolution.

As the experiment progressed to day 150, the difference in the number of mutations continued to diverge between treatments. EM1 genome in the sole presence of its host M1 (experiment 3) reached a total of 34 mutations, including 17 in a gene coding for a DNA methyltransferase and several nonsynonymous changes in genes for a DNA-binding protein, a minor tail protein, and a long-tail fiber protein, some of which may be linked to the observed decline in infectivity (Fig. 5). In contrast, EM1 genome in the experiment where all the *Sal. ruber* strains were present (experiment 4), showed 267 mutations, with a high concentration in genes related to DNA metabolism, such as a DNA primase and a single-strand DNA-binding protein. These mutations in DNA metabolism-related genes could suggest a potential disruption in viral replication, contributing to reduced infectivity (Boddin et al., 2022; Jiang et al., 2007, 2009). In the experiment containing all the bacterial strains and their viruses (experiment 5), the number of mutations in the EM1 genome increased markedly, reaching 1,331 mutations distributed across the genome. Structural and replication-related genes showed particularly high mutational loads, including 198 mutations in minor tail protein genes and over 190 combined in genes for a DNA-binding protein and a DNA primase (Supplementary Table 1). Importantly, these mutation counts represent all observed mutations above the chosen frequency threshold at each sampling point, rather than fixed mutations or strict mutation accumulation.

**Fig. 5:**
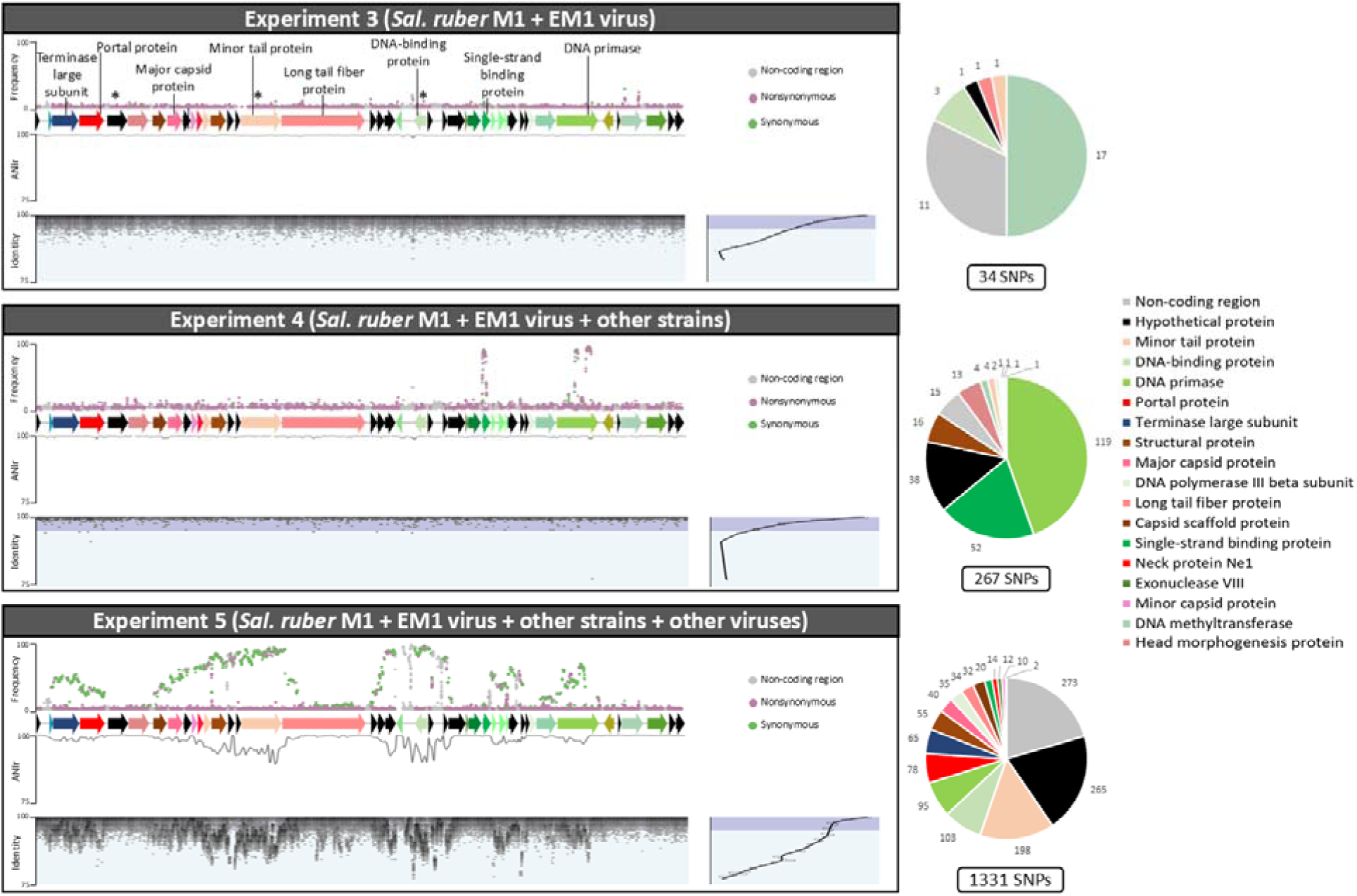
Mutations in the EM1 virus genome in the three experiments at 150 days. The three panels on the left represent experiments 3, 4, and 5, from top to bottom. In each panel’s upper section, mutations are marked as points across the EM1 genome, with the y-axis indicating mutations frequencies. Below each mutation plot is a representation of the genome of EM1, where arrows represent the open reading frames (ORFs). The middle plot depicts the ANIr (average nucleotide identity of sequence alignment reads) along the viral genome, with the y-axis showing the ANIr. In the lower part of each panel, a fragment recruitment plot is shown, where the EM1 genome is positioned at the top as the reference, and gray points represent the reads recruited, with the y-axis showing the percentage of identity for each mapped read. The histogram on the right side of this section indicates the distribution of mapped reads by identity. On the right, pie charts illustrate the protein-coding genes mutated in each experiment and the number of mutations in each ORF. Proteins selected for metabarcoding are marked with asterisks over the representation of EM1 in the first panel.

These differences were reflected in the ANI of sequence alignment reads (ANIr) along the EM1 genome (the mean nucleotide-level similarity between mapped reads and the reference genome). EM1 genome in the experiment with only the M1-EM1 pair and the condition including additional strains showed higher ANIr values (99.70% and 99.75%, respectively) indicating a more homogeneous population, compared to the more complex experiment (98.13%) on day 150 (Supplementary Table 1). The lower ANIr values observed in the presence of other viruses indicated an accelerated diversification of the viral population, with the emergence of multiple distinct genotypes or “genomovars” (defined as variants with ANI <99.5%) (Aldeguer-Riquelme et al., 2024).

The increase in genetic variability observed under complex community conditions reveals a previously overlooked role of viral coexistence in promoting diversification. Our results show that the presence of other viruses is associated with marked changes in the genetic structure of the EM1 population, both in terms of the number of mutations and their distribution across the genome (Supplementary Table 1; Fig. 5). This diversification was accompanied by a strong reduction in infectivity and a shift in host range, potentially linked to mutations in tail protein genes.

### Intracellular viral diversity exceeds extracellular detected subset

Analysis of intracellular viral populations revealed that only part of the viral diversity generated inside host cells was detected extracellularly, indicating that the extracellular fraction corresponds to a selected subset of intracellular genotypes. Three genes (marked with an asterisk in Fig. 5) were characterized by amplicon sequencing: one encoding a hypothetical protein with minimal mutations across all experiments (used as a control), and two genes highly mutated in experiment 5, encoding a minor tail protein and a DNA-binding protein, respectively. The mutation patterns in these genes were generally similar to those observed in the extracellular viral population, particularly in the condition with only the M1-EM1 pair (experiment 3) and in the condition including additional strains (experiment 4), with few differences (Supplementary Fig. 12). EM1 genome in the most complex condition, including additional strains and viruses (experiment 5), displayed a markedly higher number of mutations in the intracellular fraction in both the minor tail protein and DNA-binding protein genes (63 and 30 mutations on average across replicates) compared to experiment 3 (0 and 2 mutations) and experiment 4 (no mutations).

When comparing intracellular and extracellular fractions some differences emerged, particularly in the most complex experiment. Both the minor tail protein and DNA-binding protein regions showed a high number of mutations at nearly 100% frequency in the extracellular fraction, while the intracellular fraction exhibited more heterogeneity in mutation frequencies. These results are consistent with our earlier observations that the presence of other viruses is associated with increased genetic diversification of the EM1 population. However, based on the comparison of intracellular and extracellular data, only a subset of these genotypes appears to be detected as free virions.

### Intraspecific interactions drive host genome evolution

We also analyzed genetic mutations in the host *Sal. ruber* M1. As experiments 2, 4, and 5 included additional *Sal. ruber* strains sharing large genome regions, we focused on nine genomic regions, ranging from 1.8 to 23.5 kb, specific to the M1 strain. This strategy allowed us to unambiguously track mutations in M1 despite the presence of other closely related strains in the mixed cultures, while acknowledging that the rest of the genome could not be reliably assessed. In total, 70.1 kb were analyzed (Supplementary Table 2; Supplementary Fig. 13). Mutation counts at 150 days revealed striking differences: 9 mutations in the experiment where M1 was cultured alone (experiment 1), 224 in the experiment with other strains (experiment 2), 8 in experiment of the infection (experiment 3), 266 in the experiment of the infection plus other strains (experiment 4), and 14 in the more complex one (experiment 5), with most occurring in non-coding regions (Supplementary Fig. 14; Supplementary Table 3). Our results demonstrate that *Sal. ruber* M1 experienced more mutations in specific regions in those experiments where it evolved alongside multiple strains (experiments 2 and 4) (Supplementary Fig. 13).

These findings support the view that microbial competition plays a central role in shaping evolutionary trajectories. However, it is important to note that our analysis was limited to regions of the M1 genome that are unique and could be reliably distinguished from other coexisting strains. Therefore, although our results suggest that intraspecific competition strongly influences M1 evolution, caution is warranted in extrapolating these observations to the entire genome. Nevertheless, they underscore a critical consideration for evolutionary studies: microbial evolution and adaptation cannot be fully understood without accounting for the complex network of interspecies interactions that govern microbial life in natural environments.

## DISCUSSION

The usefulness of studying virus-host interactions within microcosms that simulate natural conditions in a stepwise manner has become increasingly evident in recent years (Blazanin & Turner, 2021; Castledine & Buckling, 2024; Chevallereau et al., 2022; Koskella et al., 2022). Understanding the ecological and evolutionary dynamics of viruses and prokaryotes in their natural environments not only enriches our knowledge of the natural world but also enables us to manipulate these communities more predictably. In this study, we demonstrate that a semi-complex biological context (encompassing *Sal. ruber* strains and their viruses) substantially influence the ecology and evolution of the *Sal. ruber* M1-EM1 pair in controlled microcosms.

At the ecological level, the findings indicated that the presence of additional *Sal. ruber* strains reduced the overall production of EM1, in good agreement with some previous studies showing that the presence of other bacterial species can lead to lower viral densities (Alseth et al., 2019; Mumford & Friman, 2017). These effects were consistently observed across different quantification methods, indicating that community composition directly impacts viral replication dynamics. This delayed population-level lysis is likely linked to the increasing abundance of coexisting strains over time, which may interfere with host-virus encounters or alter resource availability, although we deliberately refrain from attributing this effect to a specific mechanism.

In the long-term, *Sal. ruber* M1 and the EM1 virus established a stable coexistence throughout the entire experiment regardless of community complexity. This persistence was underpinned by pseudolysogeny, which we directly confirmed at the end of the experiment. These results suggest that pseudolysogeny may play a central role in maintaining long-term virus-host coexistence in nature, while simultaneously conferring resistance to the host, in line with previous studies showing that this infection mode can sustain long-term coexistence in gut environments (Schmidtke et al., 2025; Shkoporov et al., 2021). Importantly, this pattern was observed both in the simple virus-host pair and under the most complex community conditions, indicating that the establishment and maintenance of pseudolysogeny are robust to the presence of additional bacterial strains and viruses.

In the viral population, the presence of other viruses was associated with a pronounced increase in the genetic diversification of EM1, accompanied by a strong reduction in its infectivity against the original host and a shift in host range. Several studies have reported host range changes in simplified experimental evolution contexts (Bono et al., 2015; Holtzman et al., 2020; Sant et al., 2021; Subramanian et al., 2022), but only a few have explored this phenomenon under more complex conditions (Ferriol-González & Domingo-Calap, 2025; Shaer Tamar & Kishony, 2022). In multispecies settings, viruses have been shown to broaden their host range by transiently using alternative hosts as intermediates to access new hosts (De Sordi et al., 2017), which may partly explain why this host range shift occurred in the more complex experiment. Moreover, it has been observed that the presence of multiple viruses affect the host range of a given virus by driving evolutionary changes of the host that affect its susceptibility (Avrani et al., 2011; Chevallereau et al., 2022).

Beyond host range changes, the increased number and distribution of mutations observed in the presence of other viruses suggest a broader impact on viral evolutionary trajectories. While previous studies have shown that viral mutation rates can vary depending on host genotype (Duffy, 2018; Longdon et al., 2014; Sant et al., 2021), and that exposure to multiple host genotypes can promote tail gene diversification (Hernandez et al., 2024), our findings suggest that the presence of other viruses may also play an important role in shaping viral evolutionary trajectories. However, we note that these patterns may not be driven solely by changes in selective pressures, and could also reflect differences in population dynamics, such as variation in population size or genetic drift. Together, these results point to virus-virus interactions as a potentially important factor, yet still poorly understood, influencing viral evolution in complex communities.

In contrast to the viral response, the evolutionary patterns observed in the bacterial host were primarily driven by interactions with other bacterial strains. This suggests that the presence of viruses in a complex community context may constrain bacterial evolution, rather than accelerate it. This finding is striking and counterintuitive, as viruses are typically considered potent drivers of evolutionary change, and other studies showed that bacteria coevolving with multiple viruses exhibited higher mutation rates (Betts et al., 2018; Wielgoss et al., 2016). In our system, however, viral pressure may have limited evolutionary change in the host, possibly by imposing bottlenecks through lysis, constraining population dynamics, or reducing the strength of competition among bacterial strains.

The observed increase in the number of mutations under competitive conditions suggests that bacterial intraspecific competition is a strong driver of genomic evolution in the host, as suggested by previous works (Conrad et al., 2024; Goyal et al., 2022; Pharaon & Bauch, 2018). While EM1 mutations primarily emerged in contact with other viruses, *Sal. ruber* M1 genetic variability was predominantly shaped by interactions with coexisting bacterial strains. This is consistent with previous work from our laboratory demonstrating weak competitive interactions between *Sal. ruber* strains in co-culture (González-Torres et al., 2015; Peña et al., 2010). Furthermore, in a recent mesocosm experiment, we showed that strain M8 was displaced by other genotypes after being introduced at high abundance into a pond, suggesting competitive exclusion through intraspecific competition (Ramos-Barbero et al., 2024).

Overall, our results highlight the critical importance of incorporating biological complexity in studies of virus-host interactions. These findings emphasize the need to consider the multifaceted nature of microbial communities to more accurately predict virus-host dynamics and their ecological and evolutionary consequences. More broadly, they highlight that incorporating intermediate levels of biological complexity into experimental designs is essential to bridge the gap between reductionist models and natural ecosystems

## METHODS

### Bacterial strains, viruses and experimental design

*Sal. ruber* strains M1, M8, M31 and P18 were aerobically grown with gentle shaking (60 rpm) at 37°C in 25% SW (sea water, a salt solution containing the salts present in seawater at a total concentration of 25% weight/volume) with 0.2% yeast extract (Rodriguez-Valera et al., 1985). *Sal ruber* viruses used in this study, named M1EM-1, M31CC-1, M8CR-4 and M31CR41-2 (referred to here as EM1, CC1, CR4 and CR41-2, respectively, for convenience), were isolated during a previous study in our laboratory (Villamor et al., 2018).

Five experiments, with three biological replicates per experiment, were conducted in parallel (Fig. 1). In all the experiments, 25 ml of culture media were inoculated with 7.5x10^8^ cells of *Sal. ruber* M1. In experiments 2, 4, and 5, the cultures also included 2.5 × 10^7^ cells of each of the M8, M31, and P18 strains. In experiments 3, 4 and 5, 7.5x10^7^ virus-like particles (VLP) of the strictly virulent EM1 virus (multiplicity of infection = 0.01), which only infects the M1 strain, were added. Experiment 5 also contained 2.5x10^6^ VLP of each one of the CC1, CR4 and CR41-2 viruses that infect M8 and M31 strains (Fig. 1a). After inoculation, experiments were incubated for 9 days as described above, taking aliquots once a day for culture monitoring by optical density (OD) at 600 nm and additional approaches (see below). This was defined as the short-term experiment.

On day 9 (213 hours after inoculation), 250 μl of each experiment were transferred to 25 ml of fresh medium and incubated as described above. After 13 days, new aliquots were collected to quantify the bacterial and viral diversity of the cultures by microfluidics-based qPCR (see below) and transferred and incubated in the same way. This was repeated every approximately 9-13 days for 150 days. Days 9 to 150 were considered as the long-term experiment.

In addition, on days 9, 52, 84, 130 and 150, aliquots were taken for plaque forming units (PFU) quantification, host range analyses of the viruses and DNA extraction and sequencing. All experiments were conducted in triplicate.

### Flow cytometry

For each day of the short-term experiment, cells/ml and VLP/ml were quantified by flow cytometry as described in Brussaard, 2004. Briefly, 500 μl of each sample were fixed with glutaraldehyde (0.5% final concentration) at 4°C for 30 min, flash-frozen in liquid nitrogen, and stored at -80°C until use. Upon thawing, samples were diluted in Tris-EDTA buffer (pH 8), stained with SYBR^TM^Gold (0.5X final concentration), incubated for 10 min in the dark at 80°C, and cooled for 5 min at room temperature prior to analysis.

The cytometer settings were as follows: the threshold was set in blue fluorescence (300 units), FITC voltage = 500, SSC voltage = 300, forward scatter voltage = 500, and the flow rate was established as low (15 μl/min). Background noise was checked on blanks, composed by TE buffer stained with SYBR Gold. Samples were recorded with an event rate of 100-1000 events per second. Cells and VLP counts were obtained by correcting the background measured in blanks. Cells and VLP abundances were respectively expressed as cells/ml or VLP/ml.

### Plaque assay

The number of free infective EM1 viruses at each sampling point of the short- and long-term experiments was determined by plaque assay in experiments 3, 4 and 5. Aliquots of 150 μl were centrifuged at 17,000 x g for 10 min and supernatants were serially diluted in sterile SW 25%. 100 μl of dilutions 10^-4^, 10^-6^ and 10^-8^ were mixed with 500 μl of *Sal. ruber* M1 in exponential phase. Four ml of soft agar (25% SW with 0.2% yeast extract and 0.7% agar) were added and this was poured on plates with solid media. After the agar had solidified, the plates were incubated at 37 °C for 10 days until plaques were visible. The number of free infective viruses (PFU/ml) was counted and represented. On days 9, 52, 84, 130 and 150, plaque assays were also carried out to obtain individual plaques of all viruses for experiment 5, using as hosts the native strains M8, M31 and P18.

### High throughput microfluidics-based qPCR

Individual targets (i.e. the 4 strains and the 4 viruses) were quantified in each point of the short-term experiment, in both the total culture and the supernatant to quantify free viruses, using the Biomark HD high throughput microfluidics-based qPCR system (Standard Biotools, South San Francisco, USA). Primers for each target gene were designed using Geneious software 6.1.8 (Biomatters, Auckland, New Zealand) (Supplementary Table 4). From each of the experiments, 100 µl of the culture were taken for total culture analysis and 120 µl for supernatant quantification of free viruses. In the latter, a centrifugation step consisting of 10 min at 7,197 x g and 100 µl supernatant transfer to a clean eppendorf tube was performed. Both were fixed with 0.5% (final concentration) of formaldehyde for 1 hour at 4°C, diluted 10-fold in phosphate buffered saline (PBS) 1X, and stored at 4°C.

DNA targets were then pre-amplified using the PreAmp Master Mix (Standard Biotools, South San Francisco, USA) following manufacturer’s instructions. In brief, a primer pool was prepared by mixing the forward and reverse primers for all targets (Supplementary Table 4) and diluting the mixture with DNA suspension buffer (10 mM Tris/HCL and 0.1 mM EDTA pH 8) to a final concentration of 0.5 µM of each primer pair. Each sample was then pre-amplified in a total volume of 5 uL containing 1 µL PreAmp Master Mix (Standard Biotools, South San Francisco, USA), 0.5 µL of primer pool and 3.5 µL of sample. Preamplification reactions were performed on a Simpliamp Thermal Cycler (Applied Biosystems, Waltham, USA) using the following sample conditions: an initial activation cycle at 95°C for 10 min followed by 14 two-step cycles (denaturation at 95°C for 15 seconds and annealing/extension at 60°C for 4 min). The pre-amplified products were subjected to Exonuclease I cleanup (New England Biolabs, Massachusetts, USA) to remove any unincorporated primers (4 U/μL final concentration at 37°C for 30 min followed by inactivation at 80 °C for 15 min). A 1:5 dilution using DNA suspension buffer was then performed on the Exonuclease I treated samples and stored at -20°C.

Finally, pre-amplified DNA targets were quantified by microfluidics-based qPCR on 192x24 Dynamic Array^TM^ IFCs (Fluidigm, South San Francisco, USA) according to manufacturer’s instructions. In brief, 10X assay primer mixtures for each DNA target and sample pre-mixes for each sample were prepared in triplicate. For the generation of the 10X assay primer mixtures, the forward and reverse primers for a single DNA target were pooled together at a concentration of 50 uM each. 0.15 uL of this pool were then mixed with 1.35 μl of DNA suspension buffer (Tris/HCL and 0.1 mM EDTA pH 8) and 1.5 µl of 2X Assay Loading Reagent (Standard Biotools, South San Francisco, USA). Sample pre-mixes were made by combining 1.5 µl SsoFast^TM^ Evagreen® Supermix 2X (BioRad, California, USA) with 0.15 µl 20X GE Sample Loading Reagent (Standard Biotools, South San Francisco, USA) and 1.35 µl of pre-amplified DNA. The 192x24 Dynamic Array^TM^ IFCs were then primed on a Juno controller (Standard Biotools, South San Francisco, USA) and 3 uL of each 10X assay primer mix and sample pre-mix were transferred to the appropriate inlets of the IFC and loaded using again the Juno controller. After loading, the IFC was transferred to a Biomark HD instrument (Standard Biotools, South San Francisco, USA) and qPCR reactions were performed using the following cycling conditions: 95°C for 1 min, followed by 30 two-step cycles (95°C for 15 seconds and 60°C for 20 seconds) and a final melt curve analysis. Data were analyzed with the Real-Time PCR Analysis Software (Standard Biotools, South San Francisco, USA) using manually defined thresholds. All samples were run in triplicate (including the standards and negative controls). All samples, standards, and negative controls were run in triplicate, with negative controls included both in the pre-amplification and in the microfluidics-based qPCR. A total of four 192x24 Dynamic Array™ IFCs were used, each with its own standard curve consisting of five concentration points for every target, and each chip included positive controls for the eight targets individually.

### Host range through spot test and qPCR

The host range of each virus at time 0 was determined through a spot test. Briefly, 4 ml of molten 0.7% top agar of 25% SW with 0.2% yeast extract were mixed with 500 μl of bacterial cultures at exponential phase and plated on solid medium. Once solidified, 3 μl of each virus, titered at 10^10^ PFUs/ml and diluted 10^2^, 10^4^ and 10^6^ times, were added to the corresponding lawn. The spotted plates were left to dry and incubated at 37°C for 10 days. The appearance of individual lysis plaques was interpreted as evidence of productive viral infection, whereas areas with bacterial growth were attributed to resistance. This was done by exposing the 4 native strains (*Sal. ruber* strains M1, M8, M31 and P18) to the 4 viruses (EM1, CC1, CR41-2 and CR4 viruses).

The viral fractions from experiments 3, 4 and 5, at 9, 52, 84, 130 and 150 days, were obtained by centrifuging each replicate at 7,197 x g for 20 min and filtering the supernatant through a 0.22 μm filter (Millipore, Burlington, USA). For experiments 3 and 4, the host range of the EM1 virus over time was measured through a spot test against the 4 native hosts as described above.

The determination of the host range of the 4 viruses over time in experiment 5 was performed by qPCR as follows. All the plaques obtained for each native host (see plaque assay above) for each replicate on days 9, 52, 84, 130 and 150, were picked from the soft agar and resuspended in 1 ml of ultrapure water. The presence of each virus in these mixtures of plaques was checked by qPCR with specific primers (Supplementary Table 4) using SYBR Green. The experiment was carried out with the standard run in a StepOnePlus PCR System (Life Technologies, Carlsbad, USA) in a 10 μl reaction mixture with Power SYBR Green PCR Master Mix (Applied Biosystems, Waltham, USA). The reaction contained: 5 μl of 2X Master Mix, 0.2 μl of each 10 μM primer, 1 μl of sample and ultrapure water to complete volume. Conditions are detailed in Supplementary Table 5. The results were analyzed with the Applied Biosystems StepOne™ Instrument program. All samples were run in triplicate (including the standards and negative controls).

### End point colony isolation

Colonies were isolated at the final time point (150 days) by serial dilution and plating on 2% agar plates of 25% SW supplemented with 0.2% yeast extract. Plates were incubated at 37°C for up to 30 days until colonies became visible. Colonies were grown in 2 ml of culture medium at 37°C and 60 rpm. Once the cultures grew to late exponential phase, 200 μl of each were washed three times by centrifugation. DNA was extracted from washed pellets by boiling at 100°C for 10 min after resuspending the cells in 80 μl of milli-Q water. Two parallel polymerase chain reaction (PCR) amplifications were performed with each DNA, using specific primers for *Sal. ruber* M1 and the EM1 virus in parallel (Supplementary Table 4). PCR amplifications were carried out in a final volume of 25 μl, containing: 0.75 μl of 1.5 mM MgCl_2_, 2.5 μl of 10X reaction buffer, 0.5 μl of 10 mM dNTPs, 0.1 μl of Taq polymerase (5 U/μl, Invitrogen, Waltham, USA), 0.5 μl of 10 μM primers, and milli-Q water to complete the final volume. PCR conditions are described in Supplementary Table 6. PCR products were electrophoresed and UV-visualized after staining with ethidium bromide (100 μg/ml).

To assess resistance or susceptibility to the EM1 virus, colonies positives for *Sal. ruber* M1 and EM1 were exposed to the original EM1 virus and to the final virome using a spot test, as described above.

### Adsorption assay

An adsorption assay was performed in parallel with the CC1 and CR41-2 viruses and *Sal. ruber* M31 wild type. Cultures in exponential phase were diluted with fresh medium to an OD of 0.3 and 2x10^8^ cells were mixed with 2x10^6^ PFU (MOI = 0.01) of the virus. This was considered as the initial time. Mixtures were then incubated at 37°C and 60 rpm. Aliquots of 150 μl were taken along 5 h (at 1, 2, 3, 4 and 5 h), centrifuged at 17,000 x g for 8 min, and 120 μl of the cells-free supernatant were stored in ice. The numbers of PFU/ml were measured by plaque assay as described above. All the experiments were conducted in triplicate.

### Lysate experiment

An infection of *Sal. ruber* M1 with the EM1 virus at a MOI of 0.1 was carried out under the same conditions as described above. After 5 days, the infection was centrifuged at 17,000 x g for 10 minutes, and the supernatant was filtered through a 0.022 μm filter (Millipore, Burlington, USA) to remove any remaining viruses. One ml of this lysate was added to 25 ml of exponential-phase cultures of M8, M31, and P18 strains. Controls received 1 ml of fresh medium instead. Optical density was measured at 600 nm. Both the lysate-treated cultures and controls were conducted in triplicate.

### DNA extractions and sequencing

Bacterial pellets from all the experiments at 150 days were obtained by centrifugation at 17,000 x g for 10 min. The cells were washed with sterile medium, the supernatant removed, and the cell pellets stored at -80°C until extraction. DNA was extracted with the DNeasy Blood & Tissue Kit (Qiagen, Hilden, Germany) following the manufacturer’s protocol, and nucleic acids eluted in 70 μL of ultrapure water. Equal amounts of DNA from each replicate were then pooled to ensure balanced representation, capturing the overall mutational landscape of each condition.

Viruses from the same points were obtained by centrifuging at 7,197 x g for 20 min and filtering the supernatant through a filter of 0.22 μm. Dissolved DNA was removed with DNAse I from bovine pancreas (Sigma-Aldrich, St. Louis, USA) at 37°C for 30 min, with an inactivation at 75°C for 10 min. DNA was extracted with the QIAamp MinElute Virus Kit (Qiagen, Hilden, Germany) following the manufacturer’s protocol, and nucleic acids eluted in 35 μL of ultrapure water and equal amounts of DNA from each replicate were pooled. DNA from the viruses used for infecting at the initial time was also extracted as a reference.

All extracted cellular and viral DNAs were quantified using Qubit 2.0. Flourometer (Life Technologies, Carlsbad, USA), and sequenced on an Illumina Novaseq6000 (2 x 150 bp) at Macrogen (Macrogen, Seoul, South Korea).

### Metabarcoding

Three genes from the EM1 virus were selected, and specific primers targeting ∼375 bp regions were designed (Supplementary Table 7). Cell pellets were washed three times by centrifugation, and DNA was extracted and quantified as described above. Specific regions were amplified via PCR in 50 μL reactions containing: 1.5 mM MgCl , 10X buffer, 10 mM dNTPs, 5 U/μL Taq polymerase, 100 μM primers, 5 μg of the template DNA and water up to the volume. PCR conditions are detailed in Supplementary Table 8. Five ml of the amplified products were electrophoresed, stained with ethidium bromide (100 μg/mL), and visualized under UV light. The remaining product was sequenced on an Illumina Miseq (2 x 150 bp) at Fundació per al Foment de la Investigació Sanitària i Biomèdica de la Comunitat Valenciana (Fisabio, Valencia, Spain).

Mutations were identified comparing the obtained reads with the reference genomes of the viruses at initial time using Geneious software 6.1.8 (Biomatters, Auckland, New Zealand) with a minimum variant frequency of 10% and a coverage >50% to the mean coverage. Mutations were represented using the package ‘ggplot2’ v3.4.0 in R v4.2.2.

### Bioinformatics analysis

Primers and adapters were removed from sequences, and reads were filtered based on quality scores using Trimmomatic v0.36.0 (Bolger et al., 2014). Trimmed reads of initial viruses were assembled using SPAdes v3.13.1 (Bankevich et al., 2012) with the trusted option using their genomes of NCBI as a reference (GenBank accession numbers MF580955, MF580958, MF580960 and MF580960).

Mutations generated in the viruses over time were identified by comparing reads from the extracellular viromes on days 9, 84 and 150 with the reference genomes of the viruses at initial time using Geneious software 6.1.8 (Biomatters, Auckland, New Zealand) with a minimum variant frequency of 10% and a coverage >50% to the mean coverage. Mutations were represented using the package ‘ggplot2’ v3.4.0 in R v4.2.2. For the analysis of M1 sequences, the chromosomes of strains M1, M8, M31 and P18 were aligned with Mauve (Darling et al., 2004) and 9 M1-specific regions were selected where the mutations were called in the same way as for the virus.

A BLASTn was performed with the reads of each virome, using the genome of the EM1 virus as the reference. Then, the BLASTn output was filtered by best_hit option and read coverage >70%. The ANIr (average nucleotide identity of sequence alignment reads) along the genome of EM1 was calculated with a home-made script in windows of 50 bp and plotted. Recruitment plots were carried out by enveomics tool (Rodriguez-R & Konstantinidis, 2016) using enve.recplot 2 (R stadistic pluging).

### Statistical analysis

For all quantitative measurements, statistical comparisons between conditions were performed using pairwise t-tests. Each comparison was conducted using the data from the three biological replicates per condition. P-values were calculated to determine the likelihood that observed differences arose by chance, and significance thresholds are indicated in the figure legends using asterisks.

## DATA AVAILABILITY

The raw sequences used were deposited in the NCBI database with BioProject ID PRJNA1294611.

## ACKNOWLEDGEMENTS

We warmly thank the Molecular Microbial Ecology group for their valuable feedback and constructive criticism throughout this work. We also acknowledge the Genomics and Proteomics Unit of the University of Alicante for their assistance with flow cytometry. Special thanks to Antonio Sanchez-Amat and Maliheh Mehrshad for their insightful feedback and suggestions. We thank Beatriz Cámara for developing the script used in the ANIr analysis and Heather Maughan for the professional English editing and critical reading of the manuscript. This research was supported by the projects “VIRHOST” CIPROM/2021/006 (PROMETEO 2022, Generalitat Valenciana) to J.A., METACIRCLE PID2021-126114NB-C41 to F.S. and J.A. (Spanish Ministry of Science and Innovation) and CONNECTIVITY PID2024-158829NB-C41 to F.S. and J.A. (Spanish Ministry of Science and Innovation). R.S.M. received funding for his doctoral thesis from the Spanish Ministry of Science and Innovation PRE2019-087998. R.S.M., E.R.P., F.S. and J.A. are members of the National Excellence Network FAGOMA (RED2022-134837-T).

## AUTHOR CONTRIBUTIONS

F.S. and J.A. conceived the study. R.S.M., F.S., and J.A. designed the experimental approach. R.S.M. performed the experiments and analyzed the sequences under the inputs and supervision of F.S. and J.A. E.R.P. designed the primers and probes for each strain and virus. L.M.R., J.S. and M.E. provided advice and carried out the microfluidics-based qPCR. R.S.M. drafted the original manuscript. All authors contributed to manuscript final writing and approved the final paper.

