## Supplementary Information for "Biological context modulates virus-host dynamics and diversification"

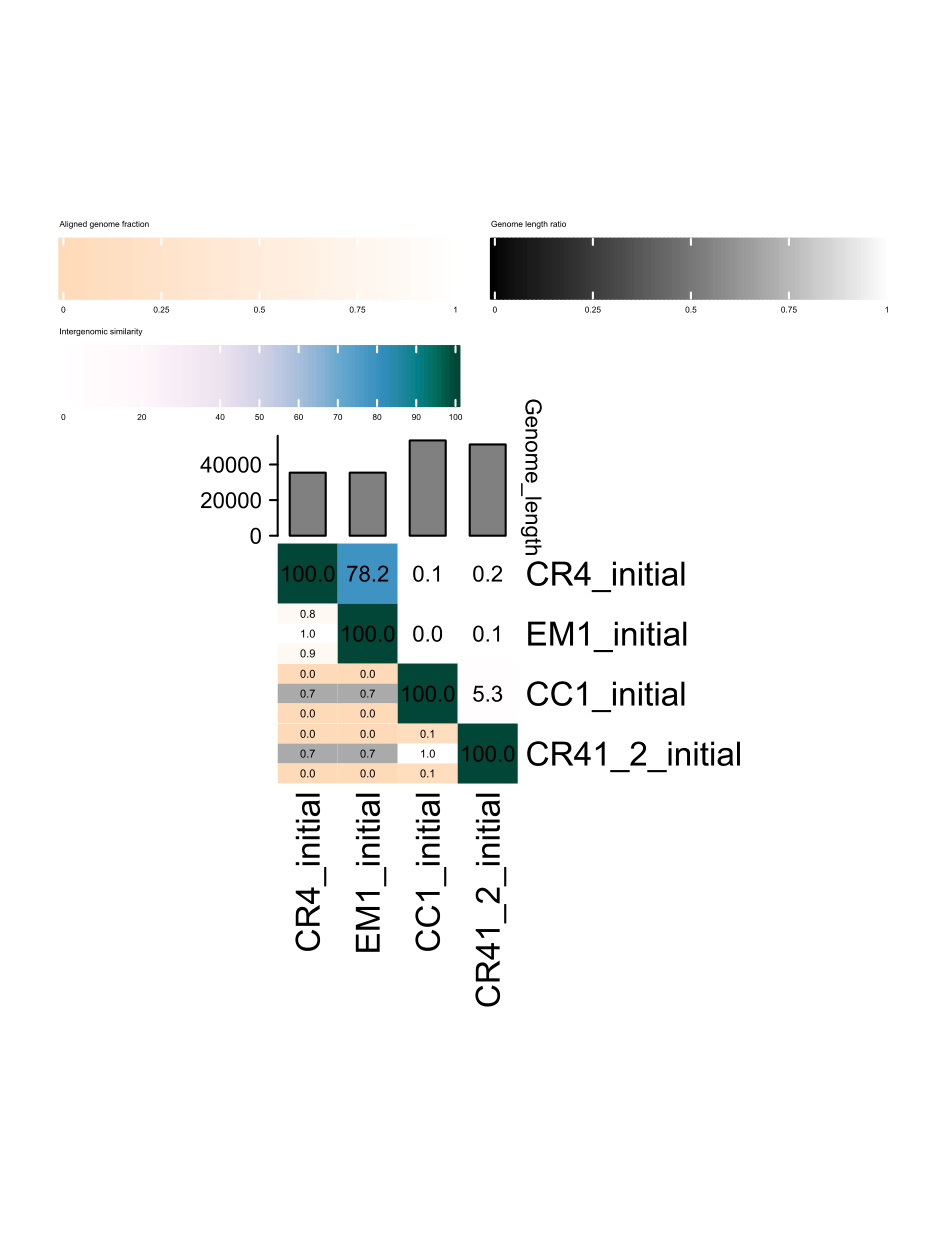
**Supplementary Fig. 1:** Intergenomic similarities of the four viruses of this study. Calculated with VIRIDIC (Moraru et al., 2020).

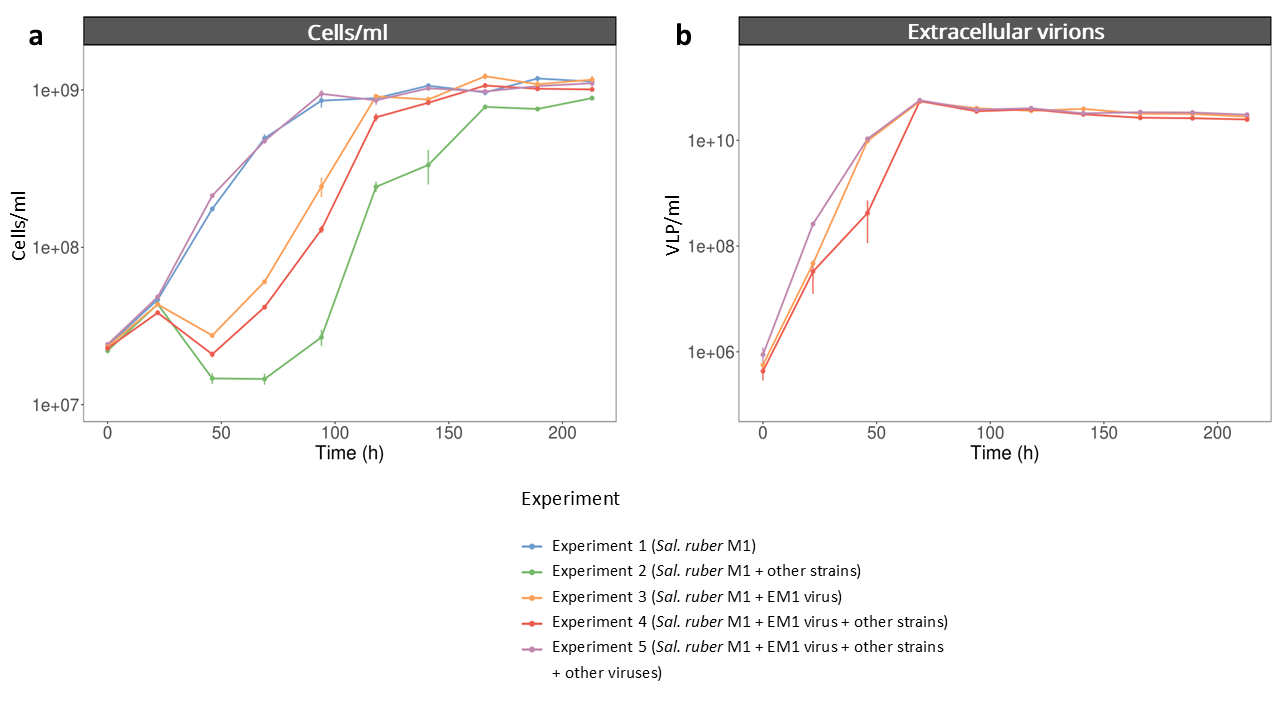
 **Supplementary Fig. 2**: **a)** Cells/ml of the five experiments measured using flow cytometry. **b)** VLP/ml of the experiments 3, 4 and 5 measured using flow cytometry. All experiments were performed in triplicate. Error bars represent the standard error of the mean across replicates.

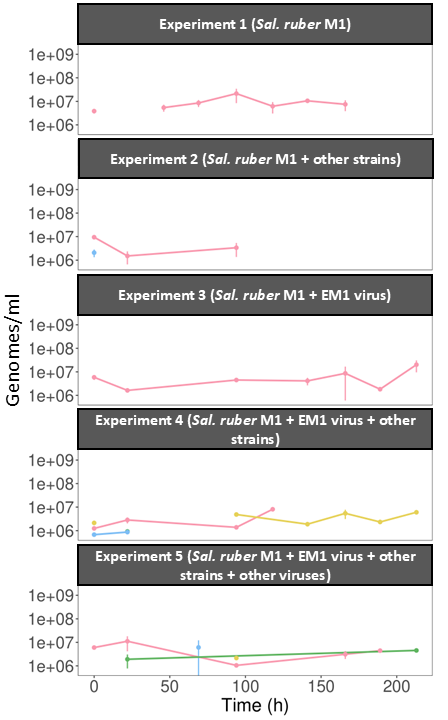

**Supplementary Fig. 3:** Genomes/ml of each strain in the supernatant in experiments 1, 2, 3, 4 and 5 (from top to bottom) measured by microfluidics-based qPCR. All experiments were performed in triplicate. Error bars represent the standard error of the biological replicates.

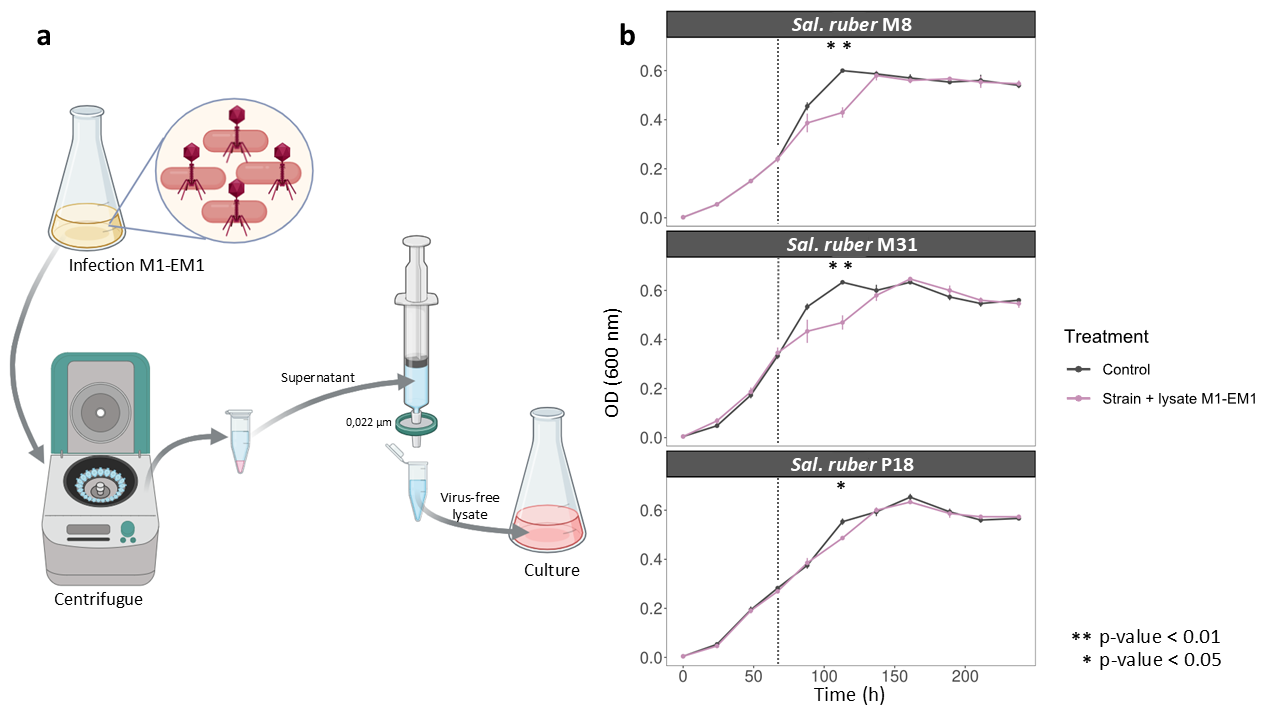
 **Supplementary Fig. 4**: **a)** The *Sal. ruber* M1-EM1 virus-free lysate was obtained by centrifugation and filtration of the *Sal. ruber* M1-EM1 culture supernatant through a 0.022 µm filter. The lysate was added at 67 hours (red dotted line) to exponential phase cultures of M8, M31 and P18 strains. **b)** Growth curves of M8, M31 and P18 amended with lysate (grey) and their corresponding unamended controls (purple). All experiments were performed in triplicate. Error bars represent the standard error of the mean across replicates. Statistical significance was assessed using pairwise t-tests between the control and the strain + lysate, with p-values indicated by asterisks. Figure created with Biorender.

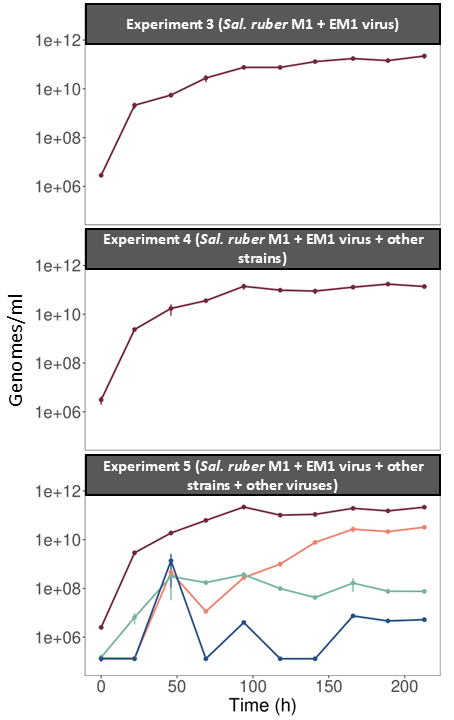

**Supplementary Fig. 5:** Genomes/ml of each virus in the whole culture in experiments 3, 4 and 5 (from top to bottom) measured by microfluidics-based qPCR. All experiments were performed in triplicate. Error bars represent the standard error of the biological replicates.

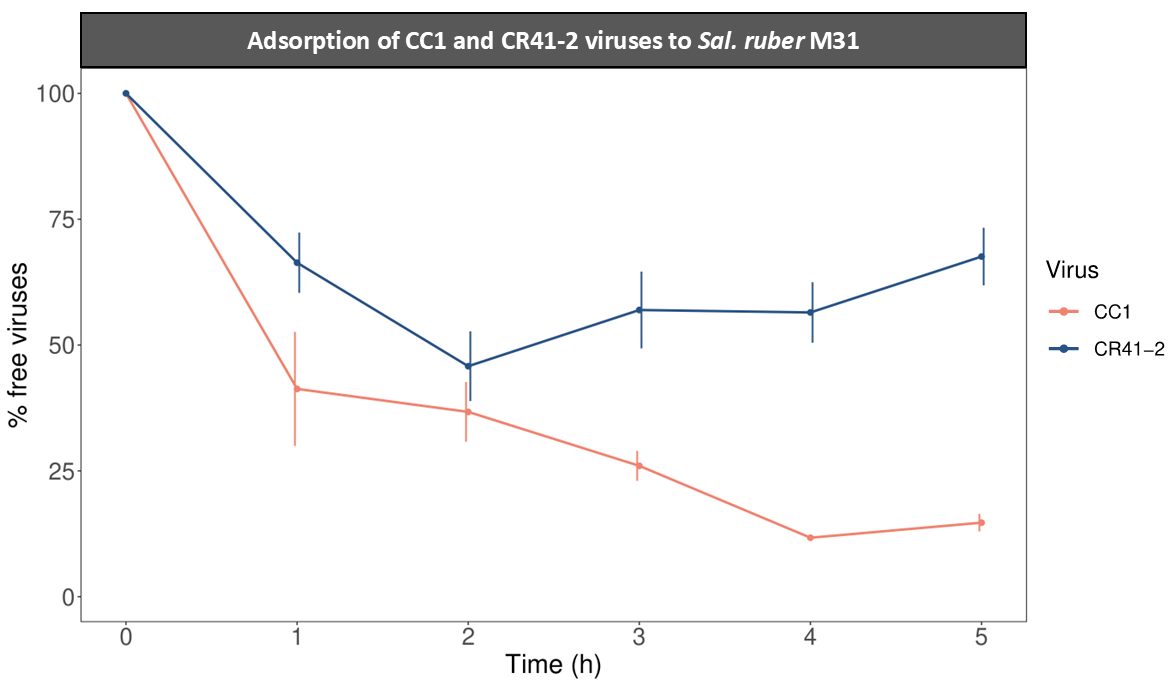
**Supplementary Fig. 6:** Adsorption experiments of CC1 and CR41-2 viruses to their M31 host. The y-axis represents the number of free viruses in the supernatant with respect to the initial viruses. All experiments were performed in triplicate. Error bars represent the standard error of the biological replicates.

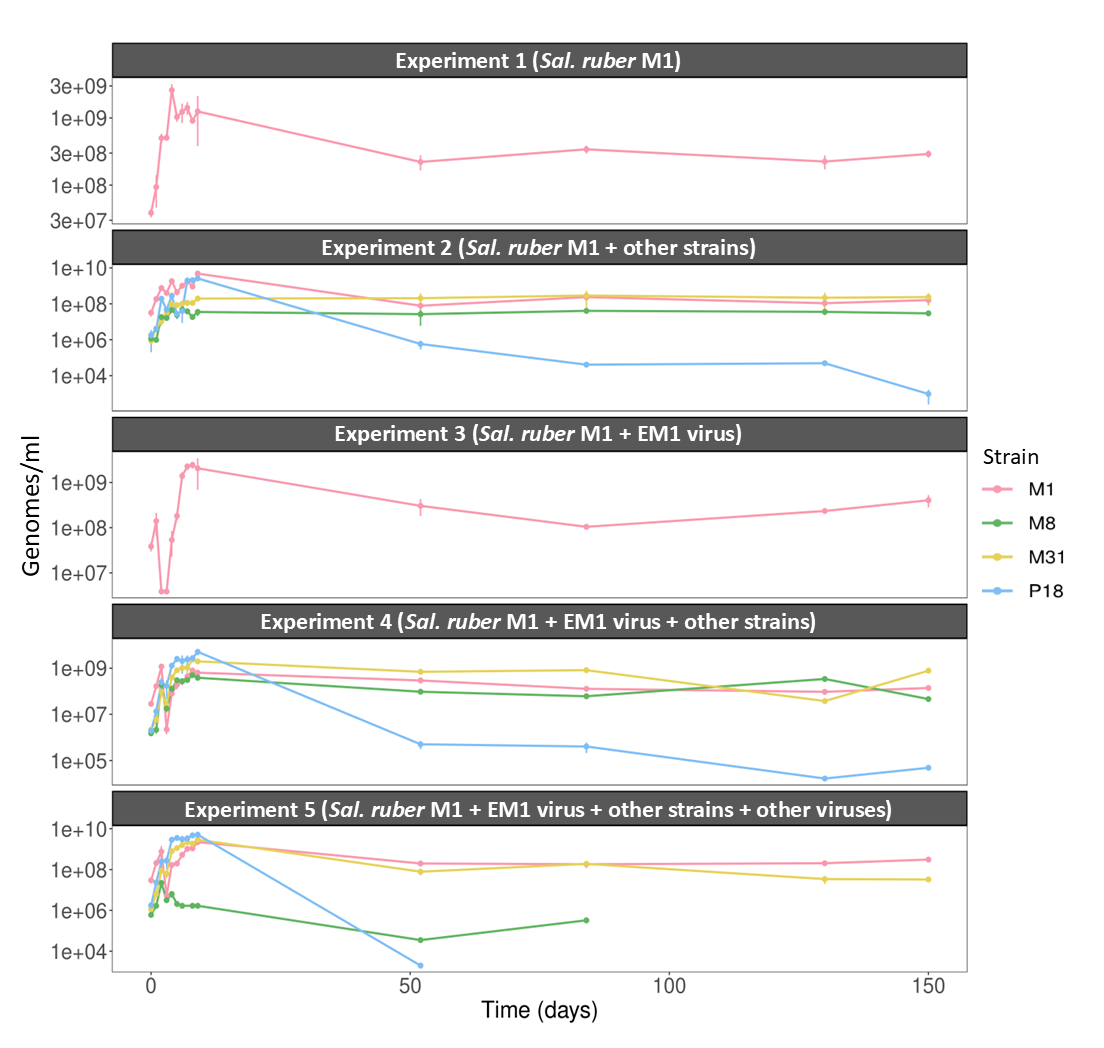
 **Supplementary Fig. 7**: Genomes/ml of each strain in the long-term in experiments 1, 2, 3, 4 and 5 (top to bottom) measured by qPCR. All experiments were performed in triplicate. Error bars represent the standard error of the mean across replicates.

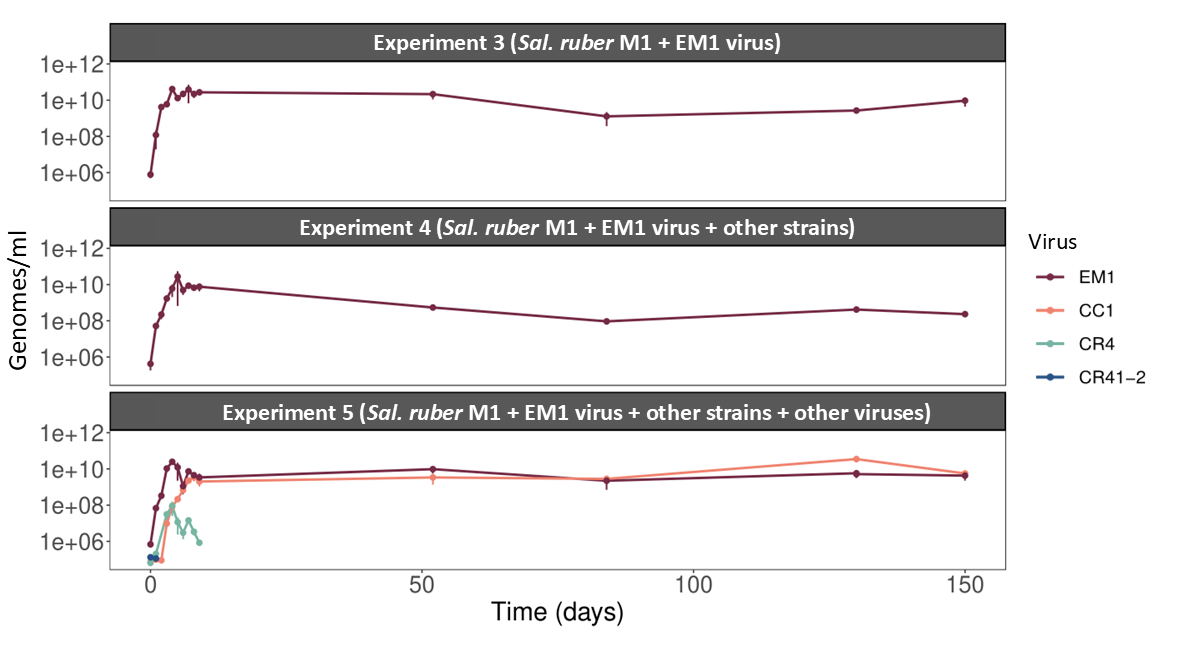
 **Supplementary Fig. 8**: Concentration of the different *Sal. ruber* viruses detected extracellularly in the long-term experiment. Genomes/ml of each virus in the long-term in experiments 3, 4 and 5 (top to bottom) measured by qPCR. All experiments were performed in triplicate. Error bars represent the standard error of the mean across replicates.

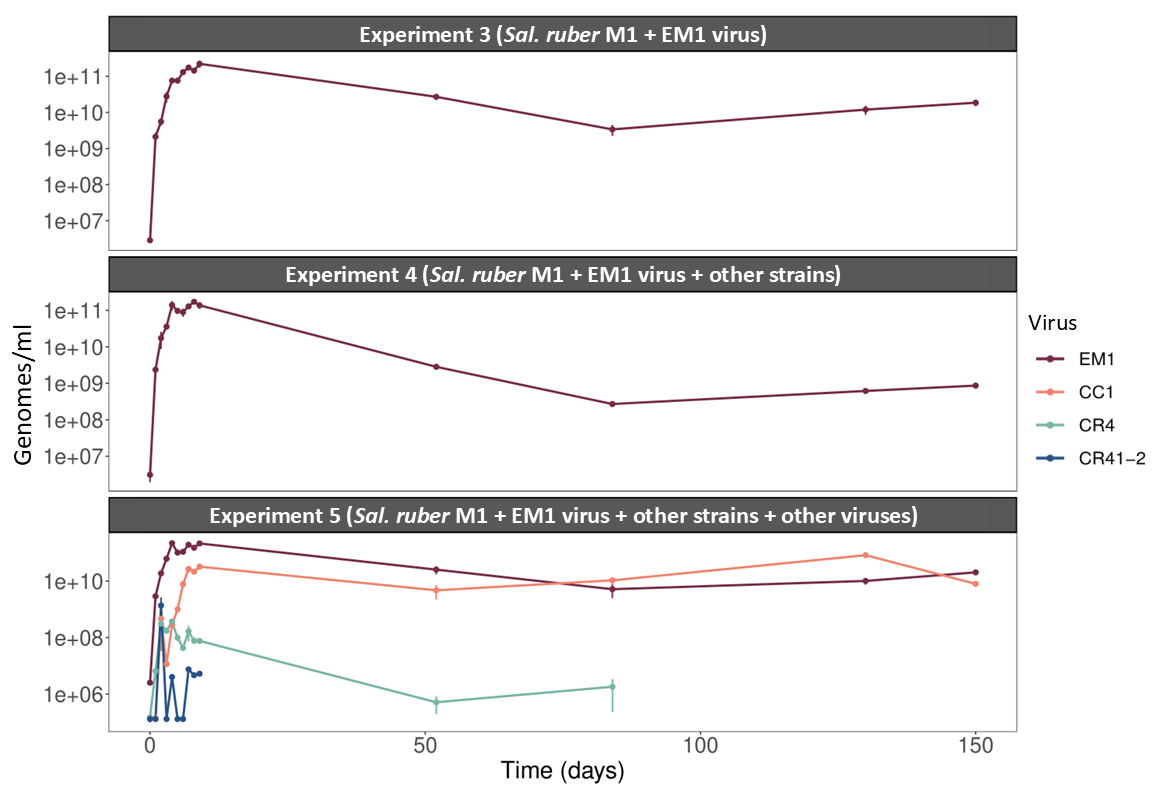
 **Supplementary Fig. 9:** Concentration of the different *Sal. ruber* viruses in the whole culture (extracellular and intracellular) in the long-term experiment. Genomes/ml of each virus in the long-term in experiments 3, 4 and 5 (top to bottom) measured by qPCR. All experiments were performed in triplicate. Error bars represent the standard error of the biological replicates.

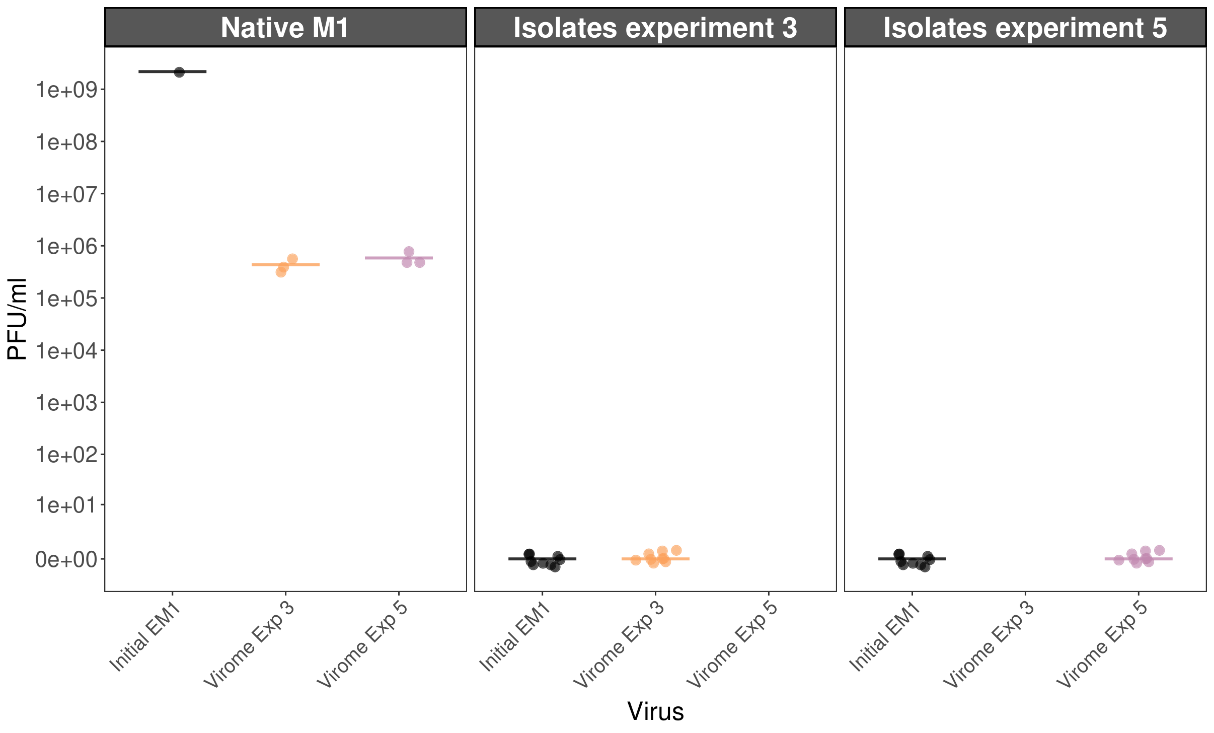
**Supplementary Fig. 10:** The susceptibility of the native M1 strain to the ancestral EM1 virus and to the final viromes from experiments 3 and 5 is shown in the first panel. The second and third panels display the susceptibility of the nine isolates obtained from experiments 3 and 5, respectively, to the ancestral EM1 virus and to their corresponding final viromes. In all panels, the mean is indicated by a horizontal bar.

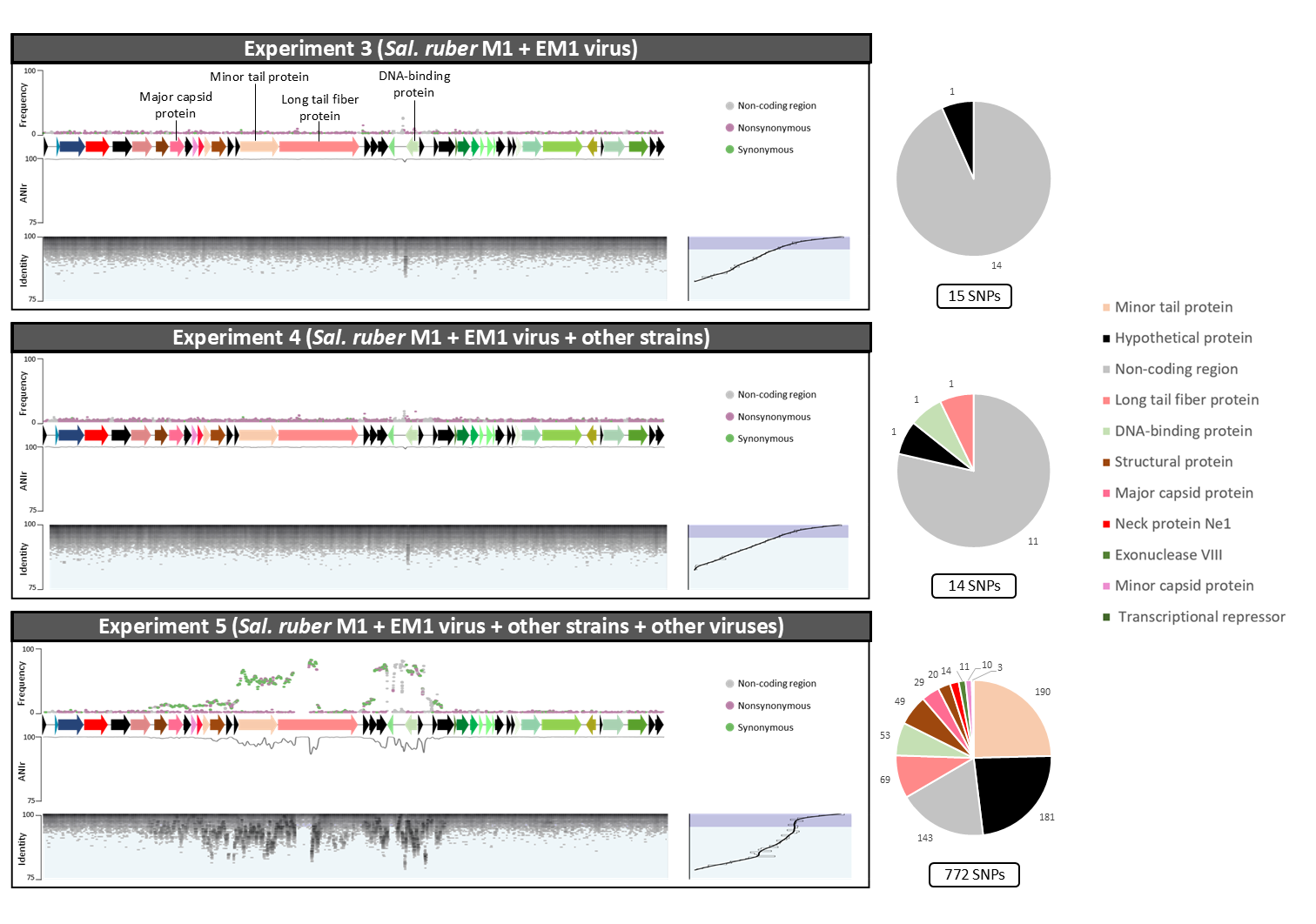
 **Supplementary Fig. 11**: Mutations in EM1 virus genome in the three experiments at 84 days. The three panels on the left represent experiments 3, 4, and 5, from top to bottom. In each panel's upper section, mutations are marked as points across the EM1 genome, with the y-axis indicating mutations frequencies. Below each mutation plot is a representation of the genome of EM1, where arrows represent the open reading frames (ORFs). The middle plot depicts the ANIr (average nucleotide identity of sequence alignment reads) along the viral genome, with the y-axis showing the ANIr. In the lower part of each panel, a fragment recruitment plot is shown, where the EM1 genome is positioned at the top as the reference, and gray points represent the reads recruited, with the y-axis showing the percentage of identity for each mapped read. The histogram on the right side of this section indicates the distribution of mapped reads by identity. On the right, pie charts illustrate the protein-coding genes mutated in each experiment and the number of mutations in each ORF. Proteins selected for metabarcoding are marked with asterisks over the representation of EM1 in the first panel.

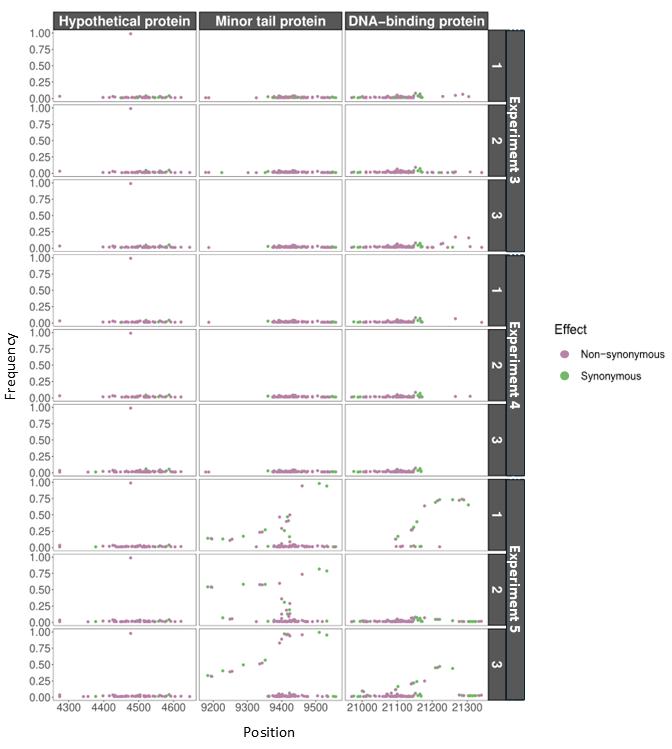
 **Supplementary Fig. 12:** Mutations in the extracellular and intracellular EM1 virus genome across the three proteins selected for metabarcoding. Each column represents one of the three selected proteins, while the different rows represent the three biological replicates of each experiment. Mutations are shown as dots along the EM1 genome, with their frequencies indicated on the y-axis.

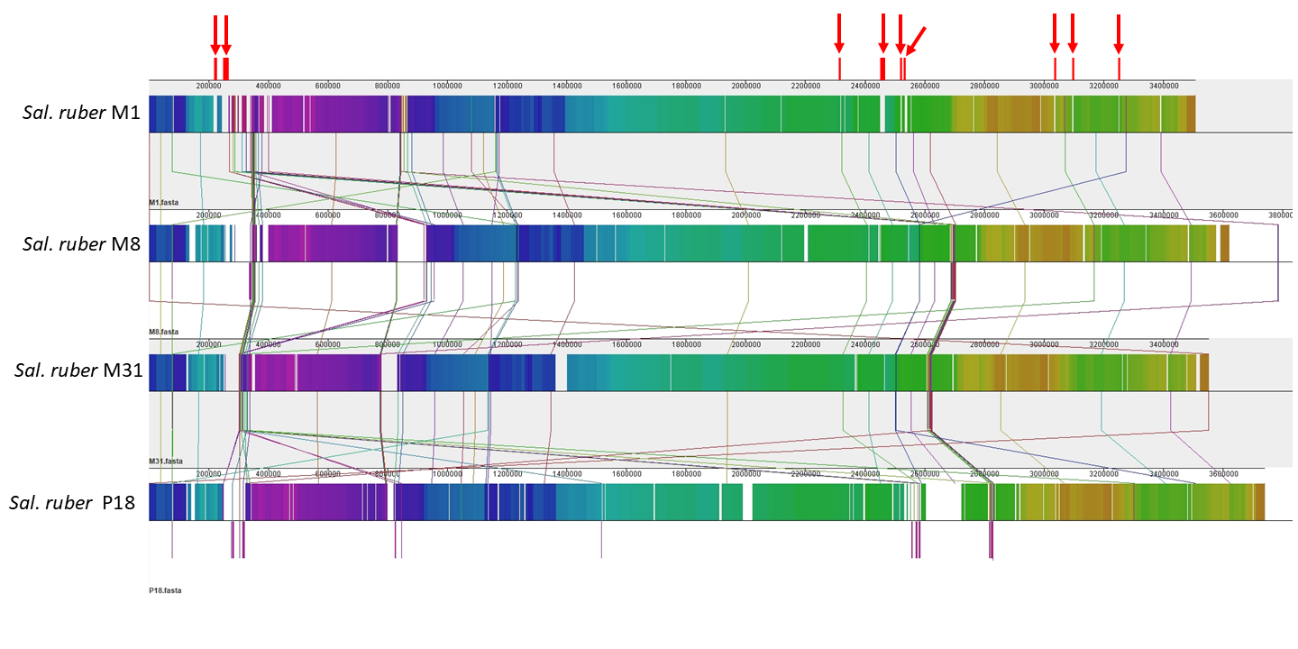
**Supplementary Fig. 13:** Alignment of strains M1, M8, M31 and P18 (from top to bottom). The genomic regions specific to M1 that were selected for mutation analysis are marked above in red. Alignment performed with Mauve (Darling et al., 2004).

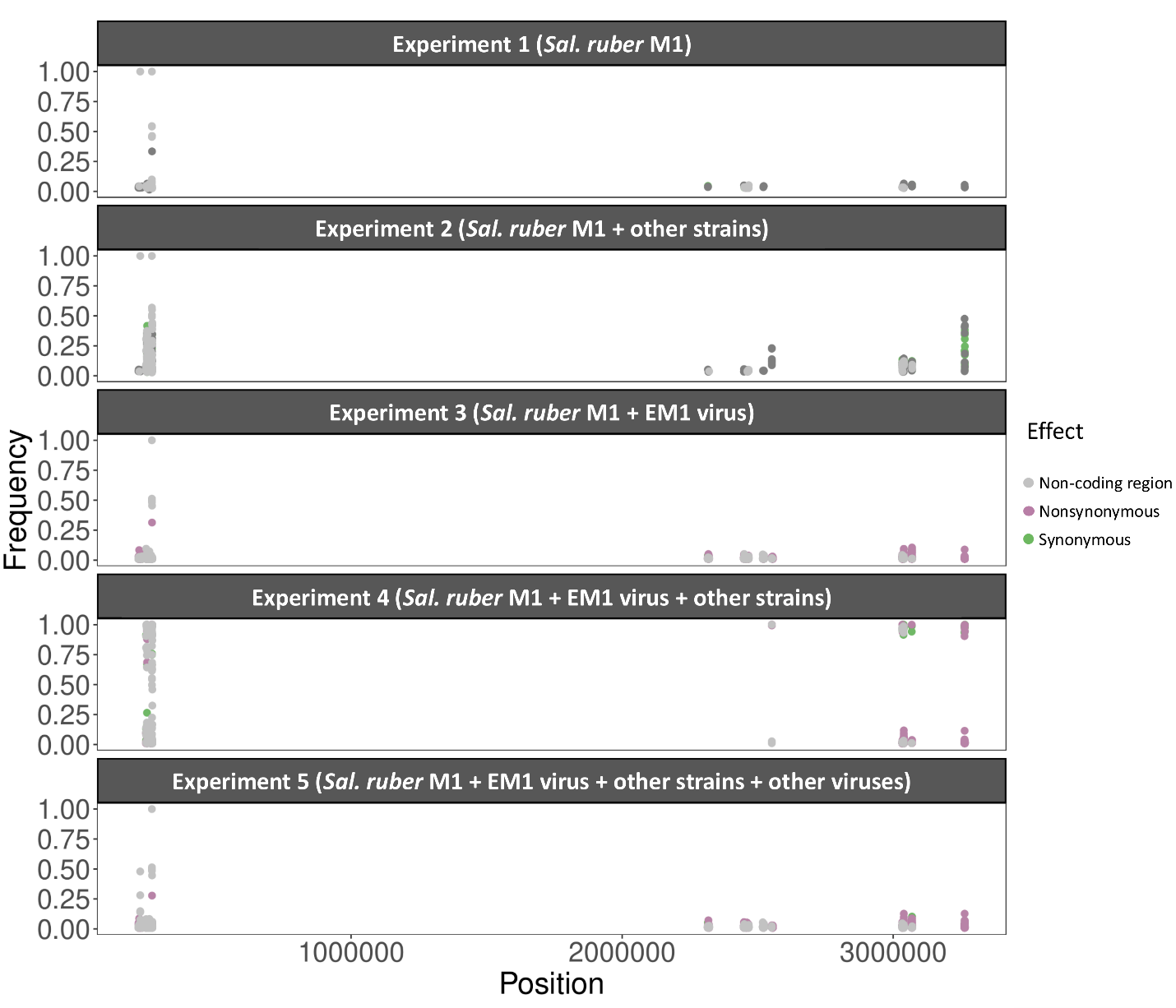
 **Supplementary Fig. 14:** Mutations accumulated in the nine strain-specific regions of the chromosome of *Sal. ruber* M1 at 150 days. The different panels represent experiments 1, 2, 3, 4, and 5, from top to bottom. Mutations are marked as points across the *Sal. ruber* M1 chromosome, with the y-axis indicating mutation frequencies.
