## Supplementary Tables for "Biological context modulates virus-host dynamics and diversification"

**Supplementary Table 1:** Number of mutations detected and ANIr values in the EM1 virus.

| **Time (days)** | **Experiment** | **SNPs (freq>10%)** | **Non-coding region** | **Nonsynonymous** | **Synonymous** | **ANIr** |
| --- | --- | --- | --- | --- | --- | --- |
| 9 | 3 | 1 | - | 1 | - | 99.85 |
|  | 4 | 1 | - | 1 | - | 99.84 |
|  | 5 | 1 | - | 1 | - | 99.81 |
| 84 | 3 | 15 | 14 | 1 | - | 99.84 |
|  | 4 | 14 | 11 | 3 | - | 99.75 |
|  | 5 | 772 | 143 | 192 | 437 | 99.22 |
| 150 | 3 | 34 | 11 | 21 | 2 | 99.70 |
|  | 4 | 267 | 15 | 168 | 84 | 99.79 |
|  | 5 | 1331 | 273 | 318 | 740 | 98.13 |

**Supplementary Table 2:** M1-specific selected regions and genes encoded withing each region.

| **Region** | **Size (kb)** | **Genes (annotation)** | **Protein ID** |
| --- | --- | --- | --- |
| 1 | 14.4 | NAD-dependent epimerase/dehydratase family protein | WP_118829866.1 |
|  |  | PIG-L deacetylase family protein | WP_118829868.1 |
|  |  | Glycosyltransferase family 4 protein | WP_118829869.1 |
|  |  | Sugar phosphate isomerase/epimerase family protein | WP_103016310.1 |
|  |  | YcaO-like family protein | WP_118829872.1 |
|  |  | Gfo/Idh/MocA family protein | WP_118829873.1 |
|  |  | HD domain-containing protein | WP_118829875.1 |
|  |  | Inositol-3-phosphate synthase | WP_118829877.1 |
|  |  | DMT family transporter | WP_118829879.1 |
|  |  | RmuC | WP_118829880.1 |
|  |  | Hypothetical protein | WP_240333595.1 |
|  |  | EamA family transporter |  |

**Supplementary Table 2 (cont.):**

| **Region** | **Size (kb)** | **Genes (annotation)** | **Protein ID** |
| --- | --- | --- | --- |
| 2 | 23.5 | Fic family protein | WP_118829892.1 |
|  |  | Type II toxin-antitoxin system VapC family toxin | WP_118829894.1 |
|  |  | nucleotidyltransferase domain-containing protein | WP_240333783.1 |
|  |  | Hypothetical protein | WP_423816085.1 |
|  |  | RES family NAD+ phosphorylase | WP_118829898.1 |
|  |  | ParS | WP_240333598.1 |
|  |  | UPF0175 family protein | WP_240333599.1 |
|  |  | Hypothetical protein | WP_240333600.1 |
|  |  | Hypothetical protein | WP_118831627.1 |
|  |  | tRNA-Asn |  |
|  |  | Hypothetical protein | WP_118829905.1 |
|  |  | Carboxypeptidase-like regulatory protein | WP_118829907.1 |
|  |  | CsgH | WP_205522017.1 |
|  |  | Hypothetical protein | WP_162899536.1 |
|  |  | tRNA-OTHER |  |
|  |  | Hypothetical protein | WP_118831628.1 |
|  |  | CsgG/HfaB family protein | WP_240333601.1 |
|  |  | Curli assembly protein CsgF | WP_118829912.1 |
|  |  | Curli production assembly/transport protein CsgE | WP_240333602.1 |
|  |  | Type II toxin-antitoxin system VapC family toxin | WP_118829894.1 |
|  |  | RNA-guided endonuclease InsQ/TnpB family protein | WP_118829914.1 |
|  |  | TnpA |  |
|  |  | Hypothetical protein | WP_262706677.1 |
|  |  | DUF6364 family protein |  |
|  |  | Nucleotidyltransferase domain-containing protein | WP_240333784.1 |
|  |  | UPF0175 family protein | WP_240331705.1 |
| 3 | 5.1 | HNH endonuclease | WP_162899668.1 |
|  |  | Tyrosine-type recombinase/integrase | WP_118831056.1 |
|  |  | Tyrosine-type recombinase/integrase | WP_118831057.1 |
|  |  | Hypothetical protein | WP_118831058.1 |
|  |  | Hypothetical protein | WP_162899670.1 |
|  |  | AAA family ATPase | WP_118831060.1 |
| 4 | 18 | Tyrosine-type recombinase/integrase | WP_118831124.1 |
|  |  | 3'-5' exonuclease | WP_118831125.1 |
|  |  | ATP-binding protein | WP_118831126.1 |
|  |  | M48 family metallopeptidase | WP_118831127.1 |
|  |  | Type I restriction endonuclease subunit R | WP_118831128.1 |
|  |  | HepT-like ribonuclease domain-containing protein | WP_118831129.1 |
|  |  | Nucleotidyltransferase family protein | WP_118831130.1 |
|  |  | Restriction endonuclease subunit S | WP_240333414.1 |
|  |  | Type I restriction-modification system subunit M | WP_118831132.1 |
|  |  | Hypothetical protein | WP_118831133.1 |
|  |  | Helix-turn-helix domain-containing protein | WP_118831134.1 |
|  |  | Tyrosine-type recombinase/integrase | WP_118831135.1 |

**Supplementary Table 2 (cont.):**

| **Region** | **Size (kb)** | **Genes (annotation)** | **Protein ID** |
| --- | --- | --- | --- |
| 5 | 4.5 | Hypothetical protein | WP_162899697.1 |
|  |  | Type I restriction endonuclease | WP_118831163.1 |
|  |  | DUF4258 domain-containing protein | WP_103017112.1 |
|  |  | Type II toxin-antitoxin system MqsA family antitoxin | WP_103017111.1 |
|  |  | Hypothetical protein | WP_103017110.1 |
|  |  | Hypothetical protein | WP_146031980.1 |
|  |  | Hypothetical protein | WP_103017108.1 |
| 6 | 3.7 | Hypothetical protein | WP_162899701.1 |
|  |  | ATP-dependent nuclease | WP_118831178.1 |
|  |  | Type IIL restriction-modification enzyme MmeI | WP_118831179.1 |
| 7 | 6.4 | Alkaline phosphatase D family protein | WP_162899727.1 |
|  |  | Alkaline phosphatase D family protein | WP_118831397.1 |
|  |  | TonB-dependent receptor | WP_272483663.1 |
|  |  | SDR family oxidoreductase | WP_118831399.1 |
| 8 | 1.9 | DUF5687 family protein | WP_118831420.1 |
| 9 | 1.9 | Prolyl oligopeptidase family serine peptidase | WP_240333519.1 |

**Supplementary Table 3:** Number of mutations detected in the genomic regions exclusive of strain M1.

| **Time (days)** | **Experiment** | **SNPs (freq>10%)** | **Non-coding region** | **Nonsynonymous** | **Synonymous** |
| --- | --- | --- | --- | --- | --- |
| 150 | 1 | 9 | 8 | 1 | - |
|  | 2 | 244 | 80 | 65 | 100 |
|  | 3 | 8 | 6 | 2 | - |
|  | 4 | 266 | 115 | 69 | 82 |
|  | 5 | 14 | 10 | 3 | 1 |

**Supplementary Table 4:** Primers used in the PCR for each target.

| ***Sal. ruber* strain / Virus** | **Primer** | **Sequence** |
| --- | --- | --- |
| *Sal. ruber* M1 | Forward | TTCGGCCTGCCTTACTCTTT |
|  | Reverse | TTTACCGTCCCCAACCAAGT |
| *Sal. ruber* M8 | Forward | TCAAGCCCCATCGGAACC |
|  | Reverse | ATCCATGCAGACGACCGG |
| *Sal. ruber* M31 | Forward | CCCGAACAGCATCACACAAT |
|  | Reverse | ATCTTTACGCGTGGCATGTC |
| *Sal. ruber* P18 | Forward | CTCGTGGTCAGACTGGTGAAT |
|  | Reverse | ACCCAGGCCGCATGTTC |
| EM1 virus | Forward | GGTCGCGGGGCTTAATATC |
|  | Reverse | CGTGTTGTTCGTTCCCCTTT |
| CC1 virus | Forward | GGTGACGGAGGGCATATTGA |
|  | Reverse | CGTTCAAGGCGTCAAGATCAT |
| CR4 virus | Forward | TGTGGTATCGCGTAGGAGGA |
|  | Reverse | GGTTTGGTAGTTGTGCGTCTTG |
| CR41-2 virus | Forward | ATTGACTCTGACGCCGCTA |
|  | Reverse | GCAATGGTACGGGTGATGATG |

**Supplementary Table 5:** qPCR conditions.

| **Nº of cycles** | **Temperature** | **Time** |
| --- | --- | --- |
| 1 cycle | 95°C | 20 seconds |
| 40 cycles | 95°C | 1 second |
|  | 60ºC | 20 seconds |
| 1 cycle | 4ºC | ∞ |

**Supplementary Table 6:** PCR conditions.

| **Nº of cycles** | **Temperature** | **Time** |
| --- | --- | --- |
| 1 cycle | 95°C | 20 seconds |
| 30 cycles | 95°C | 15 second |
|  | 59ºC | 15 seconds |
|  | 72 | 45 seconds |
| 1 cycle | 72ºC | 10 minutes |
| 1 cycle | 4ºC | ∞ |

**Supplementary Table 7:** Primers used for the metabarcoding.

| **Primer** | **Target** | **Sequence** | **Amplicon (bp)** |
| --- | --- | --- | --- |
| C_F | Hypothetical protein | TCGTCGGCAGCGTCAGATGTGTATAAGAGACAGGCTATGGATCGTCGAGGAGC | 375 |
| C_R |  | GTCTCGTGGGCTCGGAGATGTGTATAAGAGACAGCTCGTAGTTCCAGAACCCCG |  |
| MTP_F | Minor tail protein | TCGTCGGCAGCGTCAGATGTGTATAAGAGACAGGCTTGATTTCGGCCAAACGA | 378 |
| MTP_R |  | GTCTCGTGGGCTCGGAGATGTGTATAAGAGACAGGACGTCGTAGGTCGTGACAA |  |
| DBP_F | DNA-binding protein | TCGTCGGCAGCGTCAGATGTGTATAAGAGACAGCGGATTTGACGCAGAAACAGG | 377 |
| DBP_R |  | GTCTCGTGGGCTCGGAGATGTGTATAAGAGACAGCTGCACCACCAGACGTACAG |  |

**Supplementary Table 8:** PCR conditions for metabarcoding.

| **Nº of cycles** | **Temperature** | **Time** |
| --- | --- | --- |
| 1 cycle | 95°C | 3 minutes |
| 30 cycles | 95°C | 45 seconds |
|  | 51°C | 1 minute |
|  | 72°C | 2 minutes |
| 1 cycle | 72°C | 10 minutes |
| 1 cycle | 4°C | ∞ |
